# Mapping herpesvirus-driven impacts on the cellular milieu and transcriptional profile of Kaposi sarcoma in patient-derived mouse models

**DOI:** 10.1101/2024.09.27.615429

**Authors:** Xiaofan Li, Zoë Weaver Ohler, Amanda Day, Laura Bassel, Anna Grosskopf, Bahman Afsari, Takanobu Tagawa, Wendi Custer, Ralph Mangusan, Kathryn Lurain, Robert Yarchoan, Joseph Ziegelbauer, Ramya Ramaswami, Laurie T. Krug

**Affiliations:** HIV and AIDS Malignancy Branch, Center for Cancer Research, National Cancer Institute; Bethesda, MD; Center for Advanced Preclinical Research, Center for Cancer Research, National Cancer Institute; Frederick, MD

## Abstract

Kaposi sarcoma (KS) is defined by aberrant angiogenesis driven by Kaposi sarcoma herpesvirus (KSHV)-infected spindle cells with endothelial characteristics. KS research is hindered by rapid loss of KSHV infection upon explant culture of tumor cells. Here, we establish patient-derived KS xenografts (PDXs) upon orthotopic implantation of cutaneous KS biopsies in immunodeficient mice. KS tumors were maintained in 27/28 PDX until experimental endpoint, up to 272 days in the first passage of recipient mice. KSHV latency associated nuclear antigen (LANA)+ endothelial cell density increased by a mean 4.3-fold in 14/15 PDX analyzed by IHC at passage 1 compared to respective input biopsies, regardless of implantation variables and clinical features of patients. The Ki-67 proliferation marker colocalized with LANA more frequently in PDXs. Spatial transcriptome analysis revealed increased expression of viral transcripts from latent and lytic gene classes in the PDX. The expanded KSHV+ regions of the PDX maintained signature gene expression of KS tumors, with enrichment in pathways associated with angiogenesis and endothelium development. Cells with characteristics of tumor-associated fibroblasts derived from PDX were propagated for 15 passages. These fibroblast-like cells were permissive for *de novo* KSHV infection, and one lineage produced CXCL12, a cancer-promoting chemokine. Spatial analysis revealed that fibroblasts are a likely source of CXCL12 signaling to CXCR4 that was upregulated in KS regions. The reproducible expansion of KSHV-infected endothelial cells in PDX from multiple donors and recapitulation of a KS tumor gene signature supports the application of patient-derived KS mouse models for studies of pathogenesis and novel therapies.

**One Sentence Summary:** Tumor virus-driven expansion of endothelial cells with a transcriptional signature of Kaposi sarcoma in a large cohort of patient-derived xenografts provides a platform to discover cell communications within the tumor microenvironment.

## INTRODUCTION

Kaposi sarcoma herpesvirus (KSHV), also known as human gammaherpesvirus 8 (HHV8), is the etiological agent responsible for KS and four other HIV-associated diseases (malignancies or hyperproliferative conditions): primary effusion lymphoma (PEL), multicentric Castleman disease (MCD), KSHV-associated large cell lymphoma (LCL), and KSHV-associated inflammatory cytokine syndrome (KICS) (*1–5*). Kaposi sarcoma (KS) is an atypical multifocal tumor that develops in cutaneous and visceral tissues including the pulmonary and gastrointestinal tracts (*6*). Hallmark pathological features of KS are proliferative spindle-shaped cells infected with KSHV, hyper-angiogenesis, and infiltration of immune cells associated with inflammation (*7–9*). KS is classified into five epidemiological types: classic KS that typically manifests in elderly men from the Mediterranean (*10*), endemic KS in regions such as sub-Saharan Africa (*11, 12*), iatrogenic KS (*13*), KS in HIV-negative men who have sex with men (*14*), and epidemic KS associated with advanced HIV disease worldwide (*15*). Despite overlapping histological features, clinical outcomes vary significantly, ranging from an indolent course associated with cutaneous lesions in classic KS (*10*) to multiorgan involvement reported in endemic and epidemic KS (*8, 16, 17*). KS, which can also occur concurrently with MCD and/or PEL or KICS (*18*), remains a major cause of morbidity and mortality among patients living with HIV (PWH) despite adherence to antiretroviral therapy (ART) and control of HIV viremia (*19*).

Like other herpesviruses, KSHV uses the strategy of establishing a latent infection to persist for the life of the host. During latency, there is limited expression of viral products, vFLIP (open reading frame 71, *ORF71*), vCyclin (*ORF72*), latency-associated nuclear antigen, LANA (*ORF73*), and Kaposin (*K12*), in addition to KSHV miRNAs (*20–23*). LANA maintains the viral genome during cell division and the other latency factors broadly promote survival and proliferation of the infected cells (*24–28*). During productive infection following *de novo* infection of permissive cells or upon reactivation from latency, genome-wide lytic gene expression proceeds in an orderly cascade to produce viral proteins that modulate cell signaling and immune responses, in addition to proteins that replicate viral DNA and form the infectious virions (*29–33*). PEL tumors generally express KSHV latent genes. By contrast, while early studies characterized KS as primarily expressing latent KSHV genes, more recent studies have documented expression of several lytic genes including ORF75 (*34–36*). KSHV has acquired and further evolved homologs of several human genes, including a cellular cytokine (vIL-6), chemokines (vCCLs), a chemokine receptor (vGPCR), and a signaling molecule that interferes with death receptor and NF-kappa B signaling (vFLIP). Collectively, these viral proteins drive the production of growth factors and cytokines such as VEGF, PDGF and IL-6 to promote angiogenesis and tumor growth (*37, 38*). Latent and lytic factors of KSHV may thus promote angiogenesis both by direct effects in the infected endothelial cells and by indirect paracrine effects through secreted factors of viral or host origin (*8, 39*).

Among PWH with epidemic KS, antiretroviral therapy is a necessary part of treatment. For other epidemiologic subtypes and in PWH who do not have response to ART alone, chemotherapy (*6*) and the immunomodulatory agent pomalidomide are approved for KS (*40*). Pomalidomide targets host cereblon to enhance the activation of T cells and natural killer cells, as well as recognition of virus-infected cells (*40*). The response rate and the duration of the response of KS to current therapies varies depending on the agent used, the presence of concurrent KSHV-associated disorders, the extent of disease, and whether patients have received prior KS therapy (*19*). Unfortunately, despite the existing KS treatments available, some patients require intermittent KS therapy over their life span. Agents that antagonize angiogenesis, cell proliferation, and improve immune control are in clinical trials (*41–43*). Exploration of the efficacy and mechanism of action for novel interventions targeting KS pathogenesis is hampered by the inability to propagate KS-derived tumor cells in cell culture and the lack of an established pre-clinical animal model for KS.

Gammaherpesviruses that naturally infect rodents and primates have allowed for the identification of virus and host determinants of chronic infection and immune control, but KS does not arise in these model pathogen systems (*44, 45*). Xenograft models using KSHV-infected rodent precursor cells (*46, 47*), immortalized human cells (*48*), human mesenchymal stem cells (*49*) and endothelial precursor cells (*50*) lead to the development KS-like skin lesions in immunodeficient mice. More recently, a transgenic mouse line that harbors the entire KSHV genome was reported to develop angiosarcoma-like tumors with KSHV gene expression (*51*). Cell line xenograft and genetically engineered mouse models inform the processes of virus-driven angiogenesis. However, they do not fully recapitulate the heterogeneous cellular composition observed in KS tumors of patients nor do they capture the genetic diversity of clinical strains of KSHV. Patient-derived xenografts (PDXs) are generated by implanting patient tumors into immunodeficient mice, potentially providing a powerful model to capture key histological characteristics, the cellular composition and genetic variability of both the donor tumor and clinical virus isolate for *in vivo* studies (*52–55*). This facilitates mechanistic studies of virus-driven tumorigenesis and the interaction between tumor cells and the tumor microenvironment, providing a platform for the testing of novel therapies.

Herein, is the first report of the successful engraftment of cutaneous KS patient tumors in immunodeficient mice to generate KS PDX. KS was maintained in the PDX of recipient mice for long periods, up to 272 days, supporting the consistent expansion of KSHV-infected endothelial cells across multiple donors with diverse clinical features. Spatial transcriptomic analysis revealed an increase in both KSHV latent and lytic gene expression in KS PDX. The cellular complexity of the source biopsy was largely preserved in the PDX. Gene expression profiling of tumor regions defined by KS signature genes and virus infection revealed pathways that promote angiogenesis. KS PDX-derived cells with characteristics of fibroblasts were permissive for KSHV infection and produced the CXCL12 chemokine that is known to activate CXCR4 of infected endothelial cells. This newly established KS PDX model provides a powerful platform for investigating how KSHV factors promote KS tumor progression *in vivo*; paving the way for testing novel therapeutics.

## RESULTS

### KS patient xenografts maintain KSHV and retain KS histology

Thirteen participants on clinical trials with KS from diverse ethnic backgrounds and clinical histories donated skin KS biopsies for this study (Table 1). All participants had HIV-coinfection with a median CD4+ T cell count of 88 cells/µL and HIV viral load (VL) of 108 copies/ml at the time of their biopsy. Seven patients underwent KS biopsies at their initial KS diagnosis prior to systemic KS therapy with a median KSHV VL of 600 copies/10^6^ PBMCs. Six had previously received systemic KS therapy with a median KSHV VL of 1 copy/10^6^ PBMCs. In addition to KS, 7 participants also met criteria for KICS (*4*). All biopsies were taken from patients who had advanced stage (T1) KS.

**Table 1.** Clinical features of patient volunteers at time of skin Kaposi sarcoma (KS) biopsy collection.

| **Characteristics** | **Participants (n=13)** |
| --- | --- |
| **Age** (median in year, IQR) | 39.6 (34.2, 43.4) |
| **Gender** | all cisgender male |
| **Self-reported race/ethnicity, n (%)** |  |
| White | 3 (23%) |
| Black | 6 (46%) |
| Hispanic | 3 (23%) |
| Not reported | 1 (8%) |
| **HIV characteristics** | all HIV positive |
| Years from HIV diagnosis to biopsy, median (range) | 10 (0, 35.5) |
| HIV viral load, copies/mL, median (IQR) | 108 (25, 57,591) |
| CD4 T-cell count, cells/μL, median (IQR) | 88 (38, 198.5) |
| Antiretroviral therapy (ART), n (%) | 12 (92%) |
| Duration of ART in months, median (IQR) | 4.1 (0.9, 29.8) |
| **Clinical history, n (%)** |  |
| Initial KS diagnosis, no prior therapy | 7 (54%) |
| Recurrent KS diagnosis, prior systemic KS therapy | 6 (46%) |
| **KS and KS associated disease characteristics** |  |
| Years from KS diagnosis to biopsy, median (range) | 0.4 (0, 9.8) |
| KS diagnosis only, n (%) | 6 (46%) |
| KS and KSHV inflammatory cytokine syndrome, n (%) | 7 (54%) |
| Systemic KS therapy within 4 weeks of biopsy, n (%) | 3 (23%) |
| Stage of KS | T1 stage KS |
| KSHV, copies/$10^6$ cells, median (IQR) | |
| Patients with initial KS diagnosis | 600 (1, 48,571) |
| Patients with recurrent KS diagnosis | 1 (0, 1,100) |

As outlined in Fig. 1A, skin KS tumors biopsies from patient volunteers were implanted subcutaneously with extracellular matrix into NOD.Cg-*Prkdc^scid^ Il2rg^tm1Wjl^*/SzJ (NSG) or human interleukin 6 (hIL6) transgenic NOD.Cg-*Prkdc^scid^Il2rg^tm1Sug^*Tg(CMV-IL6)1-1Jic/JicTac (hIL6- NOG) immunodeficient mice within several hours of patient resection. A small portion of the KS biopsy was preserved in paraffin block for direct comparison to the matched patient-derived xenograft explanted from mice 103 d to 272 d later. Rituximab was administered to recipient mice to prevent potential onset of EBV-driven lymphoproliferation, as reported for other PDX models (*56, 57*). Among the fifteen biopsy/PDX pairs examined through hematoxylin and eosin (H&E) staining; consistent morphology was observed. A typical KS biopsy, as illustrated in Fig. 1B, revealed a histological pattern marked by variably cellular, poorly circumscribed, and infiltrative neoplastic proliferations of spindle cells intermixed with abundant dermal collagen. These proliferations formed irregularly dilated vascular channels and surrounded existing vessels. The neoplastic spindle cells ranged from flattened to plump, with amphophilic cytoplasm, minimal to mild atypia, and infrequent mitotic figures. Associated inflammatory elements included lymphocytes, plasma cells, extravasated red blood cells, and macrophages containing hemosiderin. In PDXs, the proliferating spindle cells exhibited comparable morphology to that of the original biopsy, though a higher cellular density was consistently observed in the PDX explant compared to the input biopsy (Fig. 1, B to D). The spindle cells appeared as infiltrations extending from the extracellular matrix into the surrounding tissue, in proximity to, or lining vascular structures, and as sheets of intersecting fascicles consistent with the histomorphology observed in patch, plaque or nodular stage KS. Hemosiderin-containing macrophages were occasionally present. Keratin-filled cysts were sometimes observed in the PDX when the upper epithelium was not removed from the donor biopsy prior to implant.

**Fig. 1.**
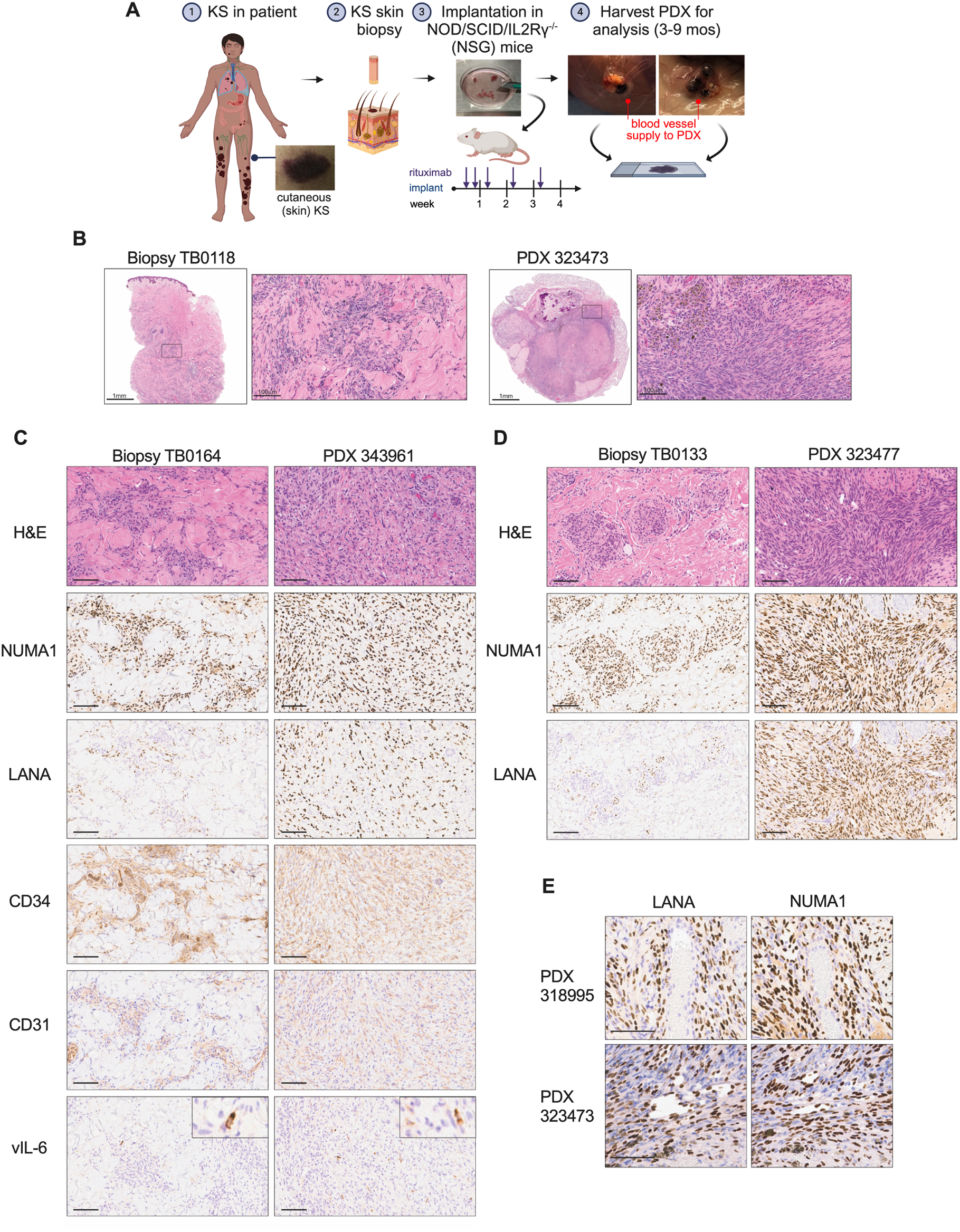
Patient biopsies implanted in mice recapitulate key features of Kaposi sarcoma. (**A**) Schematic of KS biopsy implanted into NOD/SCID/IL2Rγ^-/-^ immunocompromised mice, followed by treatment of recipient mice with rituximab. PDX tumors were harvested between 96 and 272 days. Figure prepared with Biorender. **(B**) H&E staining of a KS biopsy fragment from the patient (left panels) and a PDX derived from the implanted biopsy tissue (right panels); the area within the box in the left panel (scale bars = 1 mm) is shown at high magnification in the right panel (scale bars = 100 µm) for each tissue sample. (**C**)-(**D**) Two additional matched biopsy-PDX pairs were analyzed by H&E and then by IHC for the indicated markers for: human cells (NUMA1), KSHV+ cells (LANA), endothelial cells (CD34 and CD31), and KSHV lytic antigen (vIL-6); scale bars = 100 µm. (**E**) Two PDX tumors were analyzed for proximity of LANA+, NUMA1+ cells to vessel (top panels) and vascular-like channels (bottom panels); scale bars = 100 µm.

Immunohistochemistry (IHC) was performed on serial sections and confirmed that the proliferating spindle cells in the mouse were of human origin based on Nuclear Mitotic Apparatus Protein 1 (NUMA1) staining (Fig.1, C and D). Nuclear detection of the KSHV latency associated protein, LANA, a diagnostic criterion of KS diagnosis, was identified in all biopsies and PDX samples. The PDX spindle cells were also positive for CD34 and CD31, which are markers of endothelial cells and characterize KS spindle cells (Fig. 1C). The KSHV lytic protein, vIL-6, was detected in rare spindle cells in both biopsies and PDX (Fig. 1C). Lastly, PDX were observed to recapitulate typical features of KS including an occurrence of LANA+ cells surrounding a blood vessel (Fig. 1E, top panels) and LANA+ cells that ring vascular-like channels (Fig. 1E, bottom panels). Taken together, the KS PDXs successfully maintained the presence of KSHV and key histological features of KS tumors, and they support the expansion of LANA+ endothelial cells.

## KSHV-infected endothelial cells expand in PDX under a variety of donor characteristics and implantation conditions

KS is noted to have a variable proliferative index indicated by positivity of Ki-67 staining (*58*). To further explore increase in cellularity, we performed IHC to detect the proliferation marker Ki-67 in biopsy/PDX explant pairs. We observed an increase in Ki-67 in a representative PDX explant compared to the matched patient biopsy (Fig. 2A). Moreover, this increased detection of Ki-67 in PDX was quantifiable and consistent across 8 biopsy/PDX pairs explants, with a mean ∼4.5-fold increase (Fig. 2B). To determine if Ki-67 staining specifically indicates proliferation of LANA+ cells, co-immunofluorescence was performed (Fig. 2C). Ki-67+ cells increased from 2.5% in the biopsy to 6% of cells in the PDX, and LANA+ cells increased from 15% to 62%, while dual staining increased from 1% in the biopsy to 5.6% in the PDX (Fig. 2C). In 7 biopsy/KS pairs analyzed by IFA, the majority of LANA+ cells were not Ki-67+, however, the mean percentage of Ki-67+ in the LANA+ cells increased in the PDX (Fig. 2E). In addition, most of the Ki-67+ cells were LANA+ in the PDX compared to the paired input biopsy (Fig. 2F). Taken together, the PDX implants support proliferation of KSHV infected cells.

**Fig. 2.**
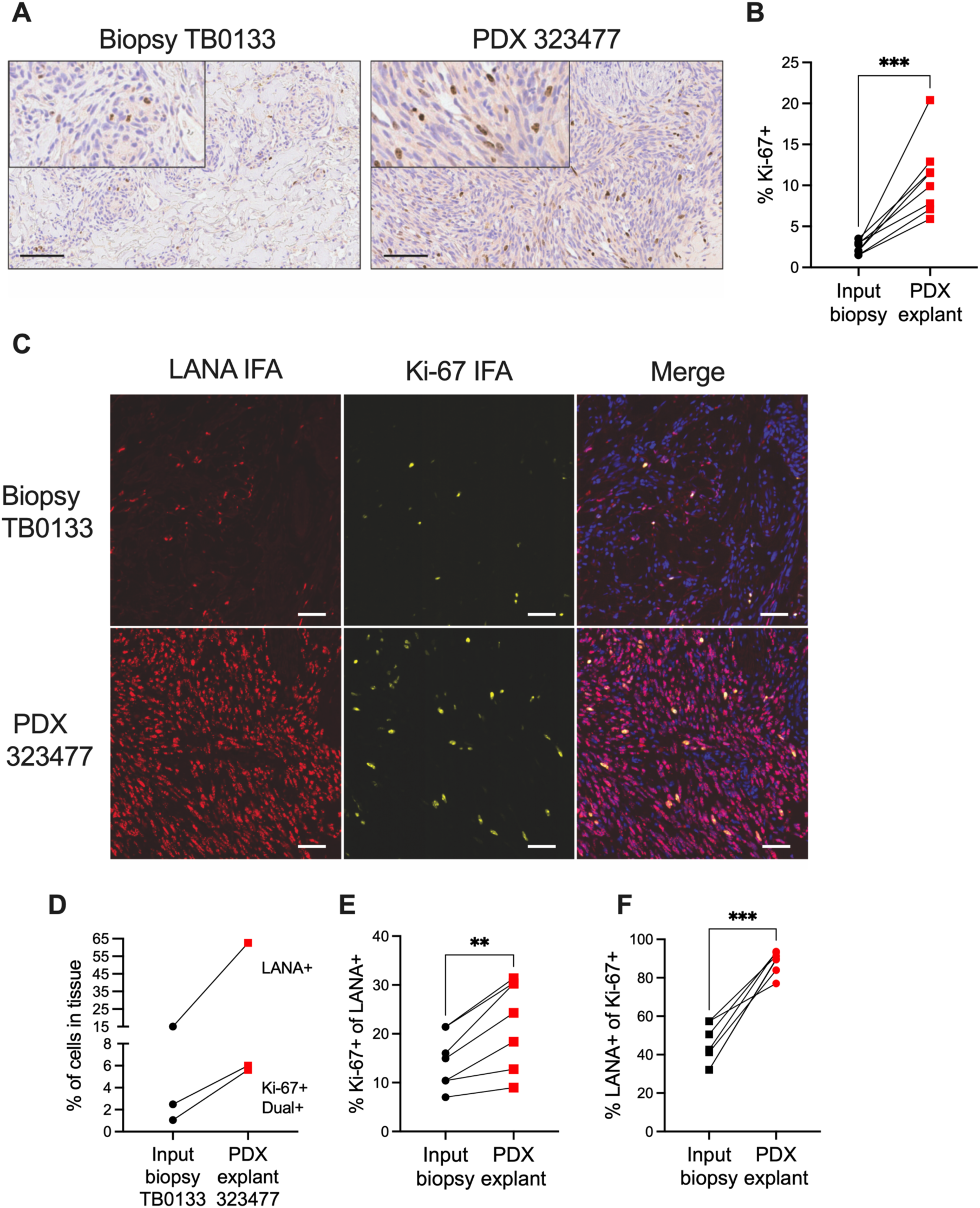
The majority of cells bearing the Ki-67 proliferation marker are LANA+ in KS PDX tumors. (**A**) IHC detection of the Ki-67 proliferation marker in a representative KS biopsy and matched PDX tumor; scale bars = 100 µm, and inset panels magnified 2X. (**B**) Percent Ki-67+ cells in IHC of biopsy/PDX pairs that were counted by automated quantification in HALO software; ***, p< 0.001, two-tailed paired t test. (**C**) IF staining for LANA (red) and Ki-67 (yellow), alone or in combination, in biopsy/PDX pair. DAPI (blue) in the merge panel; scale bars = 50 µm. (**D**) Left, quantification of IF shown in (C). (**E**), quantification of Ki-67+ relative to LANA+ cells and (**F**), quantification of LANA+ relative to Ki-67+ cells in seven additional biopsy/PDX pairs; **, p<0.01***, p< 0.001, two-tailed paired t test.

Altogether, 13 KS biopsies from patients were used as sources for 1-4 implants in separate mice, totaling 28 individual PDX (Fig. 3A). The tumor persisted in all but one PDX (27/28, 96%) and of the 17 PDX for which tissue was available 100% were positive for LANA by IHC. The remaining 11 PDX were used for passaging or cell line derivation. Since a KS-PDX system has not been previously reported, we first aimed to define the variables or combinations therein that best promote KS outgrowth (Fig. 3A). The patient donors had variable characteristics such as a new diagnosis of KS without prior systemic therapy or recurrent KS with prior therapy, and included those with KS alone or KS with concurrent KICS. We examined the impact on the density of LANA+ cells by supplementation of the extracellular matrix with vascular endothelial growth factor (VEGF), supplementation of mice with systemic hIL6, and length of implantation on the presence of LANA cells. Fold change of LANA+ cells per mm^2^ was quantified in 15 PDX at first passage (P1) compared to the corresponding input biopsy across multiple variables. This LANA+ cell density increased consistently across 14 biopsy/PDX pairs, ranging from 1.1- to 16.6-fold, a mean 4.3-fold increase (Fig. 3B). PDX 309757 had a loss of LANA+ cells to 30% of input levels. There was no difference between LANA+ cell expansion in PDX derived from patients with untreated KS versus previously treated KS (Fig. 3C) or derived from patients diagnosed with KS alone versus KS and KICS (Fig. 3D). VEGF supplementation of the extracellular matrix used for embedding did not lead to a statistical increase in the expansion of LANA+ cells (Fig. 3E). hIL6- NOG recipient mice generated serum hIL6 levels comparable to levels in patients with active KICS (Fig. 3F) but provided no benefit to LANA+ expansion in PDX (Fig. 3G). A prolonged time of implantation from 103 to 272 days did not lead to higher density of LANA+ cells in the PDX (Fig. 3H).

**Fig. 3.**
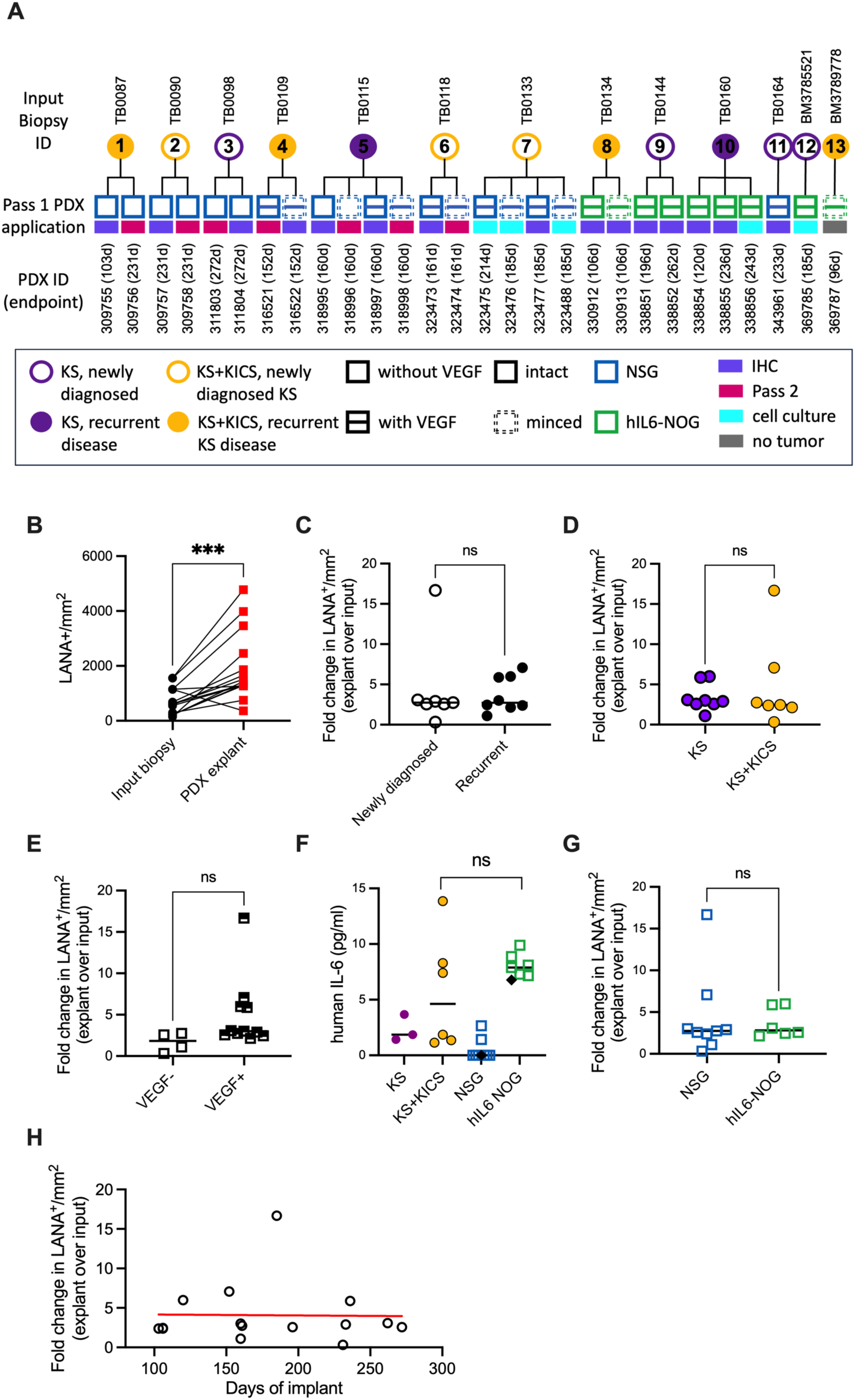
Expansion of LANA+ cells in KS PDX occurs regardless of input biopsy type or implant conditions. (**A**) Summary of all the KS biopsy (circles) and matched PDX explants (squares) described in this study, de-identified code for biopsy tissue (above circles) and recipient mouse ID (below squares) with indicated implantation times in parentheses. Clinical features of the patients (presentation of KS alone or KS in combination with KS inflammatory cytokine syndrome (KICS), primary or recurrent KS tumors) are indicated in the legend. Implantation variables included supplementation of the extracellular matrix without or with vascular endothelial growth factor (VEGF), leaving the tumor intact or minced, and implantation in NSG mice or human IL-6 transgenic NOG mice, as indicated in legend. Biopsy BM3789778 did not persist *in vivo* and not further analyzed. (**B**) Cells positive for LANA by IHC per tissue area (mm^2^) were quantified in the parent biopsy and the corresponding PDX explant; ***, p<0.001, two-tailed paired t test. (**C**)-(**E**) Fold change in LANA+ cells per tissue area of PDX explants compared to corresponding input biopsy, in patients with primary vs. recurrent KS (C), KS vs. KICS (D), and in tumors implanted without or with exogenous VEGF (E). Only two minced samples were analyzed at pass 1, precluding statistical analysis. (**F**) Levels of human IL6 in the blood of patients with KS or KS + KICS, compared to blood levels in NSG mice and hIL6-NOG mice. The black diamonds represent sera from mouse without tumor implantation. (**G**) Fold change in LANA+ cells per tissue area in PDX explants compared to corresponding input biopsy, in tumors from NSG versus hIL6-NOG mice. (**H**) Fold change in LANA+ cells per tissue area of PDX explants compared to corresponding input biopsy, by time from implant to tumor harvest.

LANA+ cell maintenance and expansion was reproducible, but the sites of tumor implantation did not change in size and there was with no evidence of ulceration or self-trauma. Next, seven P1 PDX explants were passaged again into naïve mice (Fig. 3A, Fig. S1, A and B). The tumor persisted in 5/7 (71%) these passage 2 (P2) PDX. LANA+ human cells that displayed morphological characteristics consistent with both the original biopsy and the P1 PDX were observed in 5/5 (100%) P2 tumors (Fig. S1C). Additionally, these cells were CD34+ and were observed to line blood vessels (Fig. S1D). Rare vIL-6+ cells were identified. While they did not undergo further expansion beyond the margins of the implant, KSHV LANA+ cells were maintained in this P1+P2 *in vivo* setting for up to 414 total days after biopsy, in this case 231 d for P1 (PDX 309758) followed by 183 d for P2 (PDX 316524). Taken together, with one exception, there was a consistent KSHV-driven expansion of endothelial cells in the PDX, regardless of patient baseline characteristics or implantation conditions. This supports the robustness of the skin KS PDX platform.

## Spatial analysis of KSHV-infected cell expansion and shift to lytic gene expression in KS PDX

Histopathological features, including immunohistochemical detection of KSHV LANA protein is the gold standard for the diagnosis of KS (*59*). Bulk RNA sequencing profiling has previously revealed the expression of multiple KSHV genes, including some lytic genes, in KS biopsies, in addition to *ORF73* (*34–36*). To extend beyond LANA IHC (Fig 4A, left column) and query gene expression within the tissue structure, spatial transcriptomic analysis was performed on two biopsy/PDX pairs, with an approximate 10 cells per spot resolution. These two pairs comprise biopsy samples from a newly diagnosed KS + KICS patient without prior therapy (TB0118) and a patient with recurrent KS + KICS following prior systemic therapy (TB0134). Custom probes for traditionally defined latent genes including *ORF72* and *K12* and lytic genes including *ORF75*, *K8.1* and *PAN* were spiked in with the human probe set. Viral gene expression was visualized as a heat map was overlaid on the H&E-stained section (Fig. 4A). For each biopsy, *ORF72*, *K12* and *PAN* had a larger distribution than *ORF75* and *K8.1*, which were rarely detected (Fig. 4A, first and third rows, Table S1). Analysis of the PDX for each pair revealed a dramatic expansion of the percentage of infected spots, a 3.3- and 3.5-fold increase for PDX1 and PDX2, respectively (Fig. 4A, second and fourth rows, Table S1). LANA+ regions by IHC overlapped with regions of the tumors expressing *ORF72* and *K12* and were quantitated to expand by 2.1- and 2.7-fold in the PDX of pairs 1 and 2, respectively (Fig. 4A, left column and Fig. 4B).

**Fig. 4.**
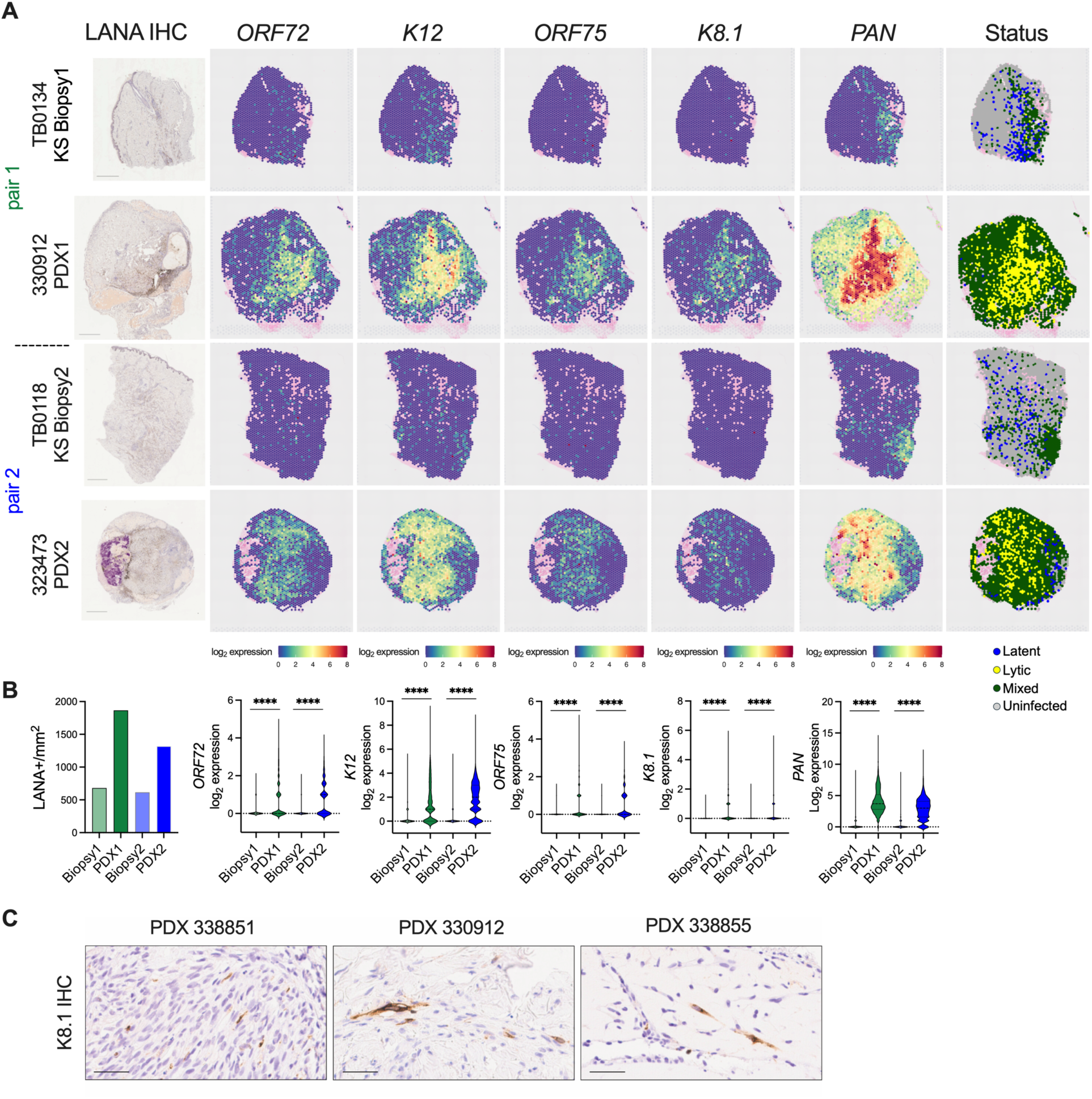
Increase in the area and intensity of KSHV gene expression in KS PDX compared to implant KS biopsy revealed by spatial transcriptome profiling. **(A)** LANA protein expression was evaluated by IHC for two matched pairs of biopsies with their corresponding PDX explants (far-left column). Adjacent sections of each corresponding tumor sample were profiled by spatial transcriptomic analysis. Spatial distribution of the indicated KSHV transcripts were evaluated by custom probes and gene expression is represented as log2 expression (calculated by read counts plus one transformed by log2) and overlaid onto the tissue using SpatialFeatures of Seurat. The far-right column portrays the status of the KSHV gene expression program across the tissues. Uninfected spots had no KSHV transcripts A latent spot was defined by the detection of at least one ORF72 or one K12 transcript, in the absence of any K8.1 or PAN transcript. A lytic spot was defined by the detection of at least one K8.1 and one PAN transcript. Mixed spots had at least one viral transcript detected but the profile fell outside the ‘Latent’ or ‘Lytic’ categories. Pink spots with low read counts were excluded from the analysis. **(B)** Left panel, cells positive for LANA by IHC per tissue area (mm^2^) were quantified for the biopsy/PDX pairs in (A) and are represented by bar graph. Right panel, log2 expression levels of the indicated KSHV transcripts are presented as violin plots for the biopsy/PDX pairs in (A); ***, p< 0.001, two-tailed unpaired t-test indicating the intensity increase of KSHV genes from a biopsy to its corresponding PDX. (**C**) IHC detecting KSHV lytic K8.1 protein was performed on PDX from three different donors.

In addition to a wider distribution of viral gene expression across the PDX, the expression levels of each viral transcript were markedly higher within the infected spots of the PDX than in the corresponding biopsy. Among the infected spots, all five genes had elevated mean log2 expression (calculated by read raw counts plus one transformed by log2) in the PDX compared to each respective input biopsy, ranging from a mean 3.8-fold increase for *K12* to a 28-fold increase for *K8.1*. (Fig. 4B). The KSHV transcript profile for each infected spot was categorized as latent (detection of at least one *ORF72* or *K12* transcript without any *K8.1* and *PAN* read), lytic (detection of at least one *K8.1* and *PAN* transcript), or mixed status (neither exclusively latent or lytic). ORF75 is considered a lytic gene based on PEL cell line patterns of gene expression but recent reports indicate high transcript levels in KS tumors (*36, 60*). Given this discrepancy, ORF75 detection was not included in these criteria for infection status. As illustrated in Fig. 4A right column, the infected spots of the biopsies were predominantly classified as latent or a mixed status, with a mean value of 37.3 or 61.9% of infected spots, respectively. The more numerous infected spots of the PDX were shifted toward a mixed or lytic status, a mean of 73.7% or 24.2% of infected spots, respectively. This increase in lytic status of virus in the PDX was validated by IHC for the K8.1 glycoprotein of the late lytic class. *K8.1* was detected in several PDX, albeit more rarely detected at the protein level than at the transcript level in PDX 330912 (Fig. 4C); K8.1 protein was not detected in matched biopsies. To summarize, our findings agree with bulk RNA sequencing reports that detect both latent and lytic gene classes in KS (*35, 36*). Importantly, this spatial approach discovered a dramatic expansion of KSHV infection within the implant and a shift towards a lytic gene expression program across the PDX tissue.

## Spatial analysis of cellular composition reveals KS tumor complexity maintained in KS PDX

Spatial transcriptomic analysis enables a more comprehensive profiling of cellular composition, as well as identifying the cell types infected with KSHV. To achieve this, the composition within each spatial transcriptome spot was estimated using the conditional autoregressive deconvolution (CARD) package using a single-cell reference of normal human skin (*61, 62*). The analysis deconvoluted the presence of eleven cell types, including immune cells (macrophages, dendritic cells, T cells, and plasma cells), endothelial cells (lymphatic and vascular endothelial cells), keratinocytes (differentiated and undifferentiated keratinocytes), stromal cells (pericytes and melanocytes) and erythrocytes (Fig. 5, A and B). The heatmap in Fig. 5A represents the proportion of different cell types in each spot and the pie chart in Fig. 5B represents the mean proportion of each cell type across all spots within each tissue. Fibroblasts comprised the largest population in both biopsies and PDXs, increasing from a mean proportion of 42% in all spots in the biopsies to 61% in PDXs (Fig. 5B). Consistent with the expansion of CD31+ or CD34+ cells by IHC (Fig. 1C), the mean proportion of endothelial cells increased from a mean 17% in biopsies to 24% in PDXs (Fig. 5B). Keratinocytes were primarily detected in the epidermis of the original biopsies with their proportion in PDXs dropping from a mean of 15% in the original biopsies to 3.5% in PDXs. In addition, a decrease in the mean proportion of immune cells was observed in the PDXs, from 7.9% in biopsies to 5.1% in PDXs (Fig. 5B). Overall, the tumor cellular complexity of the source biopsies was maintained for an extended period in mice until experimental endpoint, 106 d and 161 d for PDX1 and PDX2, respectively.

**Fig. 5.**
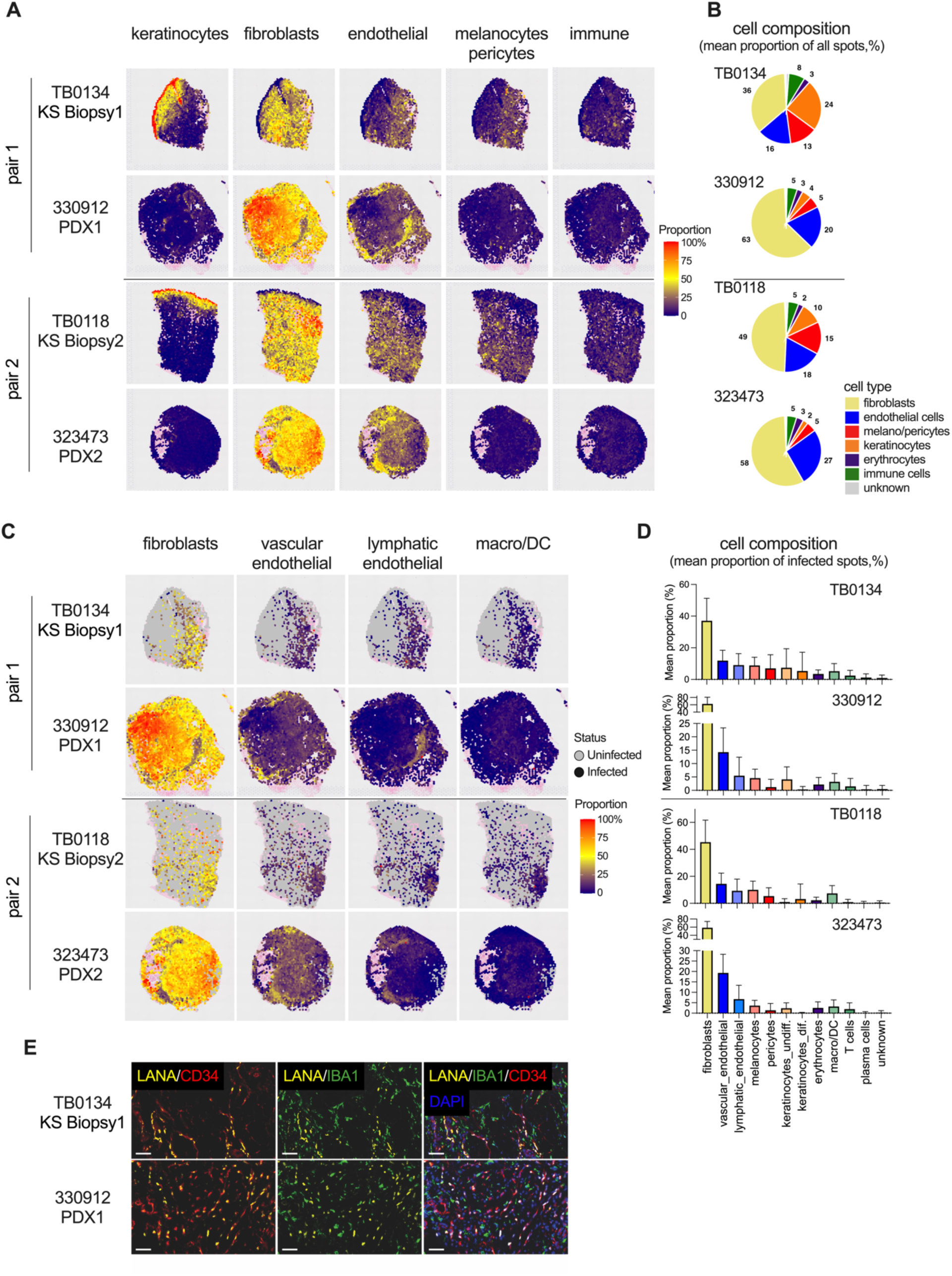
The cellular composition is broadly conserved in KS PDX compared to KS biopsy. **A)-D)** Two matched pairs of biopsies were analyzed using the ‘conditional autoregressive deconvolution’ (CARD) package of Seurat using skin single-cell reference data. **(A)** The spatial distribution of indicated cell types is presented as a heat map overlay that indicates the relative proportion of the indicated cell type per spot. **(B)** Pie charts summarize the mean percentage composition of cell-types across all spots in each tissue in (A). **(C)** A finer but selected spatial distribution of indicated cell types as performed in (A) only for infected spots, i.e., at least one KSHV transcript (*ORF72, K12, ORF75, K8.1,* or *PAN*) detected. **(D)**. Bar graphs summarize the mean proportion with standard deviation bar of each cell-type in the infected spots of each tissue in (C). **(E)** IF of KSHV LANA to examine co-localization with cells bearing the endothelial cell marker CD34 (left panel) or the macrophage marker IBA1 (middle panel) in the indicated biopsy/PDX pair. DAPI (blue) in the merge panel; scale bars = 50 µm.

KSHV infects multiple cell types in culture. In KS tissue sections, KSHV LANA is detected in cells that bear markers of endothelial cells and fibroblasts/mesenchymal stem cells (*63, 64*) and in sections adjacent to those that stain for macrophages (*65*). When examining the KSHV-infected spots, fibroblasts comprised the highest mean proportion of 41% and 60% in the biopsies and PDXs, respectively. The mean proportion of vascular endothelial cells increased from 13% in biopsies to 17% in PDXs and lymphatic endothelial cells decreased slightly from 9% in biopsies to 6% in PDXs (Fig. 5, C and D). Macrophages comprised a mean 6.2% proportion of all the infected spots in the biopsies and a mean 3.2% proportion in the PDXs (Fig. 5, C and D). The mean proportion of pericytes and keratinocytes was reduced in infected spots of PDXs (Fig. 5D). Cells in a biopsy/PDX pair were examined at the single-cell level via immunofluorescence assay staining for KSHV LANA, the endothelial cell marker CD34, and the macrophage marker IBA1 (*66*). LANA+ cells co-stained with CD34, but not IBA1, in both biopsy and PDX. Notably, macrophages demonstrated proximity to KSHV-infected cells both in biopsy and PDX (Fig. 5E), consistent with immune cell infiltration noted in KS tumors (*67*). Taken together, KSHV infected regions of the tumor are not homogenous. Multiple cell types including immune cells are interspersed within the infected regions of the biopsy, and that cellular complexity is maintained as virus infection expands in the PDX.

## Gene expression profile of clusters with KS signature genes and regions of infection in KS and KS PDX

We next used a clustering-based comparison of gene expression across integrated samples, independent of viral transcripts, to circumvent the bias driven by extreme differences in viral gene expression between biopsies and PDX samples. Samples were hierarchically integrated, normalized and clustered with Seurat. In the scType package, the mixed cellular composition is represented by scType scores, wherein the predominant cell-type signature for each cluster is assigned based on the highest scType score (Fig. 6A, Table S2). ‘Unknown’ reflects a cluster for which no cell type predominated or could not be assigned using the references provided.. In addition, a custom cell-type assignment termed ‘KS signature’ was generated from differentially expressed genes (DEGs) based on published bulk RNA sequencing datasets of KS samples (*35, 36*) that were further refined to identify the 5 top predictors of KSHV+ spots in the biopsy/PDX tissues: *ADAMTS4, ADAM19, FLT4, STC1, CLEC4M* (Fig. 6A, Table S2).

**Fig. 6.**
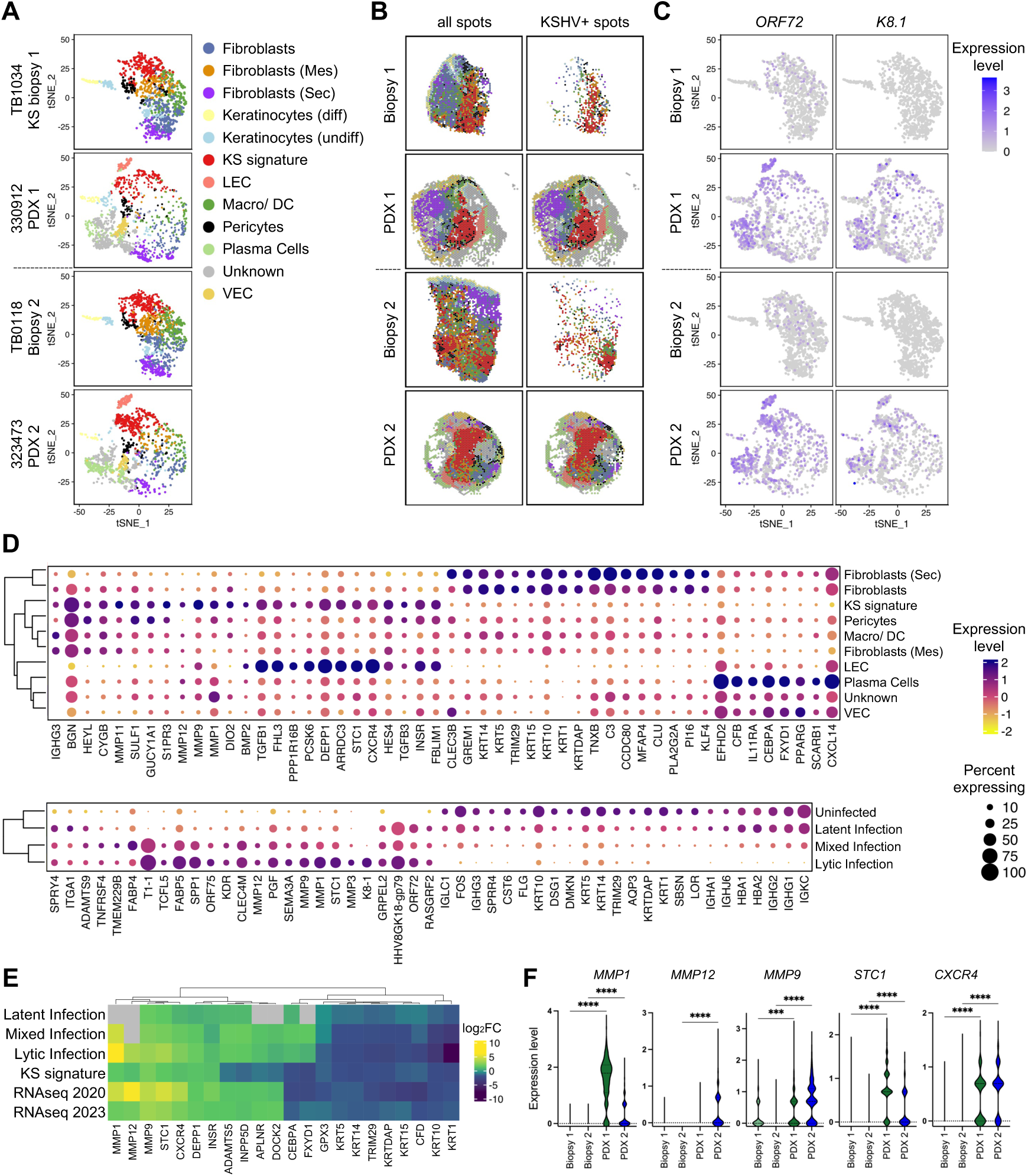
Differentially expressed genes in clusters with KS gene signatures overlap with differentially expressed genes in KSHV infected spots in KS biopsies and KS PDX. **(A)** Automated cell type identification based on marker genes for 12 cell types was performed using scType and overlayed on the tSNE plot. (**B)** Spatial distribution of cell types across samples visualizing all spots (left panel) or infected spots with at least one KSHV transcript (*ORF72, K12, ORF75, K8.1,* or *PAN*) detected (right panel). **(C)** tSNE plot of SCT normalized expression of a marker of latent (ORF72) or lytic (K8.1) KSHV infection per sample. (**D)** Expression of differentially expressed genes (DEGs) in the KS signature cluster compared to the other clusters (upper panel) and DEGs in spots with the indicated infection status (as described in Fig. 4A, right panel) compared to the uninfected population (lower panel). The top 25 up- and downregulated genes (adj p-value <0.001) in the KS signature cluster (top panel) and top 15 up- and downregulated genes (adj p-valye <0.001) in KSHV infected populations (lower panel) were chosen for visualization; duplicated genes are treated the same as single hits. SCT transformed expression levels are indicated by the scale bar and the percentage of spots with detectable gene expression per cluster are shown by circle size. Keratinocytes were excluded from the analysis to prevent bias towards strong keratinocyte markers. (**E)** Expression of DEGs found in the KS signature cluster, infected clusters and published bulk RNA sequencing datasets comparing KS to normal skin was summarized in a heatmap;RNAseq datasets published in 2020 (*35*) and 2023 (*36*), KS signature and infected spots reported in (D). (**F)** Violin plots of the SCT normalized expression for the upregulated genes in (E) in the KS signature clusters of each sample. ***, p< 0.001; ****, p< 0.0001.

When the clusters were spatially overlaid onto the tissues (Fig. 6B), these cell assignments from Seurat largely overlapped with the distribution of cells determined by CARD analysis (Fig. 5, A and B). Keratinocytes and secretory fibroblasts were more distal around the borders of the tumors, with VECs more apparent in the periphery of the PDX. Regions of the tumor identified as the KS signature were adjacent to regions dominated by LECs in the PDX and interspersed with regions dominated by mesenchymal fibroblasts, pericytes, macrophages, and plasma cells in the biopsies and the PDX. Since each spot within a cluster represents multiple cells, other cell types with the next ranked scType scores were evaluated. Notable, in one of the two clusters identified as KS signature, clusters identified as LEC and VECs were assigned the second and third highest scType scores, respectively, while the LEC cluster displayed high scType scores for KS signature and VEC markers (Fig. S2A). The spatial proximity of clusters based on the 5 KS signature genes to LECs is consistent with the overlap of marker genes between the clusters as expected from the known predominance of endothelial cells in KS.

Regions of virus infection, based on the detection of at least one KSHV transcript as described in Fig. 5C, were dominated by LECs, KS signature, plasma cells, VECs and the unknown cluster (Fig. 6B, right panel and Fig. S2B). We next examined the viral gene expression across clusters of the tSNE plots. The latent *ORF72* transcript was detected at low levels in multiple clusters of the biopsy and was most highly expressed in the LECs and unknown cluster of the PDXs, followed by KS signature and plasma cell clusters (Fig. 6C). The lytic K8.1 transcript was detected in very few spots of the biopsies but followed a similar pattern of distribution in the PDX clusters (Fig. 6C).

To further delineate the specific gene profile distinguishing clusters of specific cell-types, DEGs for each of the annotated cell types were determined. Analysis of DEGs for the KS signature cluster compared to other cell-type clusters identified 103 genes with a two-fold change in expression; the top twenty-five upregulated and twenty-five downregulated genes are shown as a bubble plot that indicates both the mean expression level and the percentage of spots in a cluster that express the indicated DEG (Fig. 6D, upper panel). The DEGs for the KS signature cluster were further analyzed for enriched gene ontology (GO) biological processes, and the linkage of genes and biological processes were visualized using network plots (Fig. S3). Upregulated genes of the KS signature cluster encode products involved in organizing the extracellular matrix that include *MMP9*, *MMP1*, *MMP11* and *MMP12*, the development and differentiation of endothelial cells that include *STC1*, *S1PR3* and *CXCR4*, and the bone morphogenetic protein and Notch signaling pathways that include *BMP2*, *PCSK6*, *HEYL* and *SIPR3* (Fig. 6D and Fig. S3A). The top downregulated gene in KS signature clusters encode proteins engaged in lipid localization and transport, *PLA2G2A*, *C3*, *PPARG* and *CLU,* and negative regulators of cell growth, *PI16* and *GREM1* (Fig. 6D and Fig. S3B). Notably, most of the top regulated genes in KS signature clusters demonstrated similar expression profiles in LEC (Fig. 6D). Both clusters have a high proportion of viral transcript detection (Fig. S2B), suggesting DEGs are, in part, being driven by KSHV infection.

Next, 448 DEGs specifically regulated in KSHV infected clusters that encompass latent, lytic and mixed spots (Fig. 4A) compared to uninfected clusters were analyzed (Fig. 6D, lower panel). The top upregulated genes in KSHV lytic and mixed infection spots are predicted to encode proteins that regulate angiogenesis including *ADAMTS9*, *KDR* and *PGF*, endothelial cell development including *KDR* and *STC1,* and cell-substrate adhesion, *SPRY4*, *ITGA1*, *ADAMTS9*, *KDR*, *MMP12* (Fig. 6D and Fig. S3C). In KSHV lytic and mixed infection spots that predominant in the PDX, the top downregulated genes encode proteins involved in epidermis development, leukocyte mediated immunity, and cellular response to external stimulus that include *FOS* and *AQP3* (Fig. 6D and Fig. S3D). This is consistent with the sharp reduction of keratinocytes (Fig. 5A and Fig. 6A) and the use of rituximab for B cell ablation in the PDX (Fig. 1A). Notably, DEGs enriched in pathways that regulate angiogenesis, endothelial cell development and cell-substrate adhesion, such as *CLEC4M*, *MMP9*, *STC1*, *ADAMTS9 and SPRY4,* were also identified in latent infected spots, albeit expression levels were lower than in lytic and mixed clusters (Fig. 6D and Fig. S3C). Immune cell response networks were reduced in the infected clusters (Fig. S3D).

Those 103 and 448 DEGs enriched in the respective KS signature and KSHV infected clusters based on spatial analysis were next compared to the DEGs identified from published bulk RNA sequencing datasets from endemic and HIV-associated KS tissues (*35, 36*). As described in the Venn diagram of Fig. S3E, 22 genes were characterized as overlapping DEGs between the four DEG lists. The majority of these DEGs shared a similar trend of upregulation or downregulation across all datasets (Fig. 6E). The expression level of these overlapping DEGs were further analyzed by infection status (Fig. 6E). Broadly, there was a gradient of increased expression of the upregulated DEGs moving from latent to mixed to lytic infection status, with a similar pattern for downregulated DEGs. The relative expression levels of the top 5 upregulated DEGs, *MMP12*, *STC1*, *MMP1*, *MMP9* and *CXCR4*, were analyzed across the biopsy/PDX tissues. DEG expression was on average higher in the PDXs compared to their respective biopsies (Fig. 6F).

Taken together, the KS signature gene expression profile was maintained in PDX and behaved similarly to the LEC cluster that was spatially adjacent in the PDX. The PDX also had a notable expansion of a mixed and lytic infection status that drives DEGs in pathways that promote endothelial cell development and angiogenesis. Thus, these findings from the spatial analyses of the biopsy/PDX pairs largely agree with those previously reported in bulk RNA sequencing analysis of KS tumors. Taken together, the PDX provides a robust platform to examine virus-driven changes in the endothelial cells of KS tumors.

## Fibroblast-like cells derived from KS xenograft produce CXCL12 upon KSHV infection

The PDXs maintained KSHV infection for extended times in the mice, but it was not clear if cells derived from the xenograft would maintain or support infection. Cells freshly isolated from KS tumors have limited doubling times prior to reaching senescence (*68*) and rapidly lose the virus even if initially infected (*69*). To reduce the potential for reactivation of latently infected cells that might harbor KSHV in the process of explant culture, we adopted a strategy reported for the derivation of EBV+ nasopharyngeal (NPC) PDX cell lines. The Rho kinase inhibitor Y-27632 suppresses tetradecanoyl phorbol acetate (TPA)-induced EBV lytic replication in cell culture and blocks spontaneous virus reactivation in EBV+ NPC PDX cells (*70, 71*). To investigate the impact of Y-27632 on KSHV, iSLK-Bac16 cells were induced to reactivate in the presence of 10 uM Y- 27632 and monitored for productive replication. Late lytic gene expression was reduced by the Rho kinase inhibitor at 72 hpi in both qRT-PCR and immunoblot analysis, with a three-fold reduction in infectious particle production (Fig. S4).

To derive cells from KS PDX for cell culture propagation, two KS PDX explants were embedded in extracellular matrix and cultured in media supplemented with the Rho kinase inhibitor, Y-27632 (Fig. 7A). Cells migrated out from the extracellular matrix plug and reached confluence over a 45 d period and were then propagated for fourteen passages prior to reaching senescence. The two KS PDX xenograft (KSX) derived cell cultures that were developed, KSX-476 and KSX-488, stained positive for the human cell marker NUMA1 (Fig. S5A) and matched the source KS donor through short tandem repeat analysis (Table S1). KSX-476 and KSX-488 expressed tumor associated fibroblast (TAF) markers including platelet derived growth factor receptors (PDGFR)- α and -β, α-smooth muscle actin (α-SMA), fibroblast activation protein (FAP) and S100A4 (*72–74*), with some variance noted between the two cell cultures (Fig. 7B). Minimal levels of the endothelial markers CD31 and vascular endothelial growth factor receptor (VEGFR)-3 were detected on KSXs (Fig. 7B, S5B to E).

**Fig. 7.**
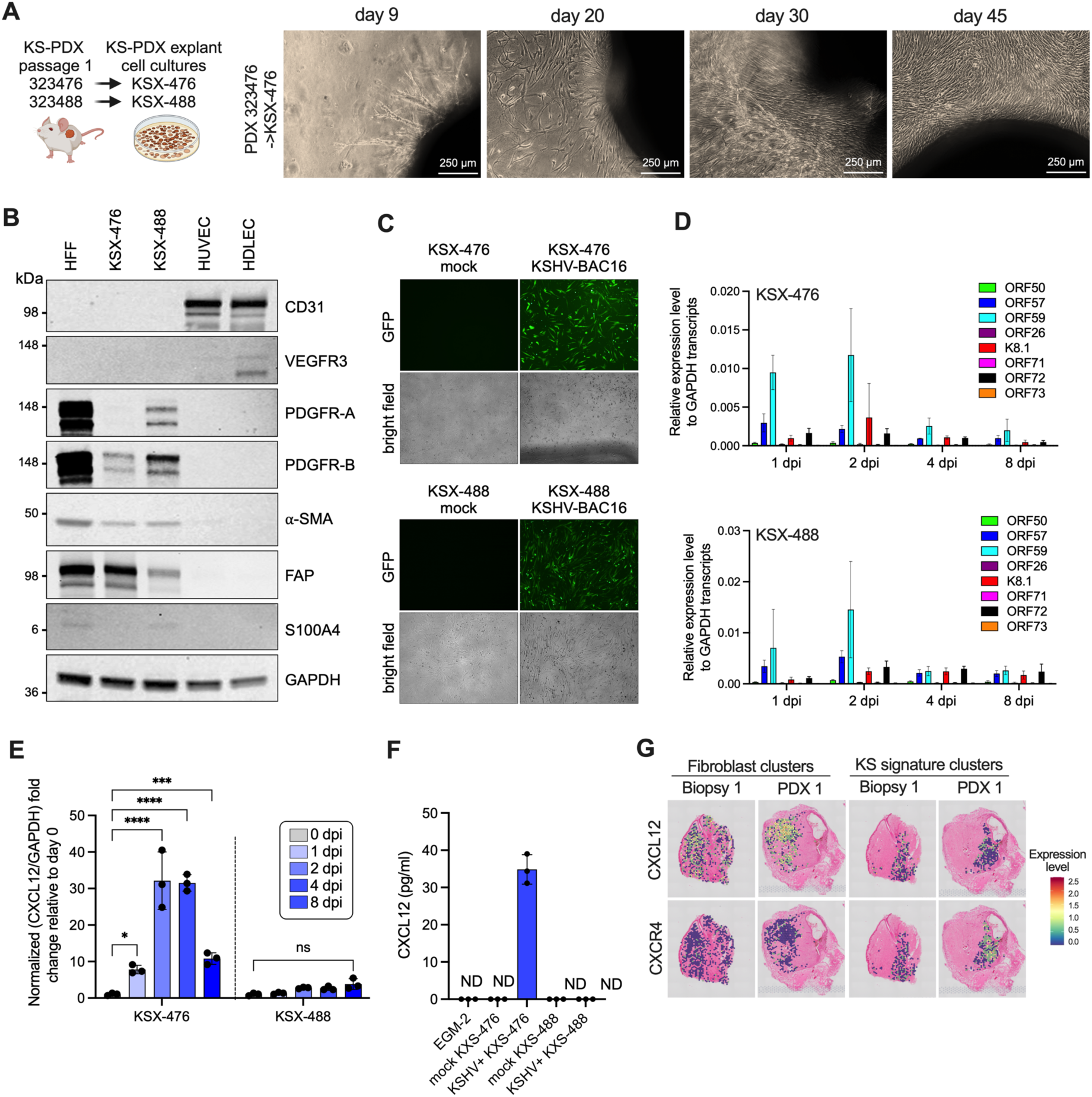
KS PDX-derived cells bear markers of tumor associated fibroblasts that produce CXCL12 upon KSHV infection. **(A)** Two KS PDX explants were embedded in extracellular matrix in the presence of ROCK inhibitor; cells sprouted and expanded over 45 d to generate PDX-derived cells (KSX-476 and KSX-488) that were maintained for 15 passages. **(B)** Cell lysates of KSXs, HFF and endothelial cells were analyzed for markers of endothelial cells (CD31 & VEGFR3), fibroblasts (PDGFR-α & PDGRF-β) and TAFs (a-SMA, FAB, and S100A4) by immunoblot. **(C)** KSX-476 and -488 were inoculated with KSHV-BAC16 at MOI 1.0, and KSHV infection was monitored by the GFP+ cells by microscopy. **(D)** KSX-476 and -488 were infected with KSHV-BAC16 at MOI 1, and the level of the indicated KSHV transcripts was determined relative to GAPDH over 8 d. **(E)** The fold change in GAPDH-normalized expression of CXCL12 in KSHV- infected KSX-476 and -488 was determined relative to day 0 prior to infection. **(F)** The conditioned media collected from mock and KSHV-infected KSXs at 4 dpi was subjected to ELISA to measure the extracellular level of CXCL12 protein. **(G)** Spatial distribution of CXCL12 (upper) and CXCR4 (lower) normalized expression levels in fibroblast and KS signature clusters in the indicated samples.

The KSHV genome was detected only in early passages of KSX-488 prior to reaching a level below the limit of detection (Fig. S5F). However, both KSXs were permissive for *de novo* KSHV infection with KSHV-BAC16 (Fig. 7C), comparable to the levels of infection in other permissive cell types, human foreskin fibroblasts and primary LECs (Fig. S5G). Expression of KSHV lytic genes *ORF50*, *ORF57*, *ORF59*, *ORF26* and *K8.1* peaked at 2 dpi and declined over the 8 d time course in both KSXs; whereas expression of latency associated genes *ORF71*, *ORF72* and *ORF73* was minimal and largely unchanged over the time course (Fig. 7D). There was no detectable release of infectious virions in the conditioned medium of the infected KSXs (Fig. S5H). Taken together, cells derived from the KS PDX explants share features of TAFs and are permissive for *de novo* KSHV infection.

We next explored whether viral infection of these PDX-derived stromal cells may influence the tumor microenvironment of KS. CXCR4 was a top upregulated gene in infected regions of KS biopsies and PDX (Fig. 6D to F) and is upregulated in KSHV-infected endothelial cells in culture (*64*). CXCL12 expression is elevated in multiple tumor types and engages receptors CXCR4 and CXCR7 to promote proliferation and survival of cancer cells (*75*) and are detected in KS tumors (*76*). A qRT-PCR screen examining genes associated with angiogenesis identified CXCL12 as a virus-responsive gene in KSX-476 cells (Fig. S5I). CXCL12 transcripts were significantly upregulated in KSX-476 but not KSX-488 upon infection (Fig. 7E). Importantly CXCL12 protein was produced and released from KSX-476 at 4 dpi (Fig. 7F). Spatial analysis of KS/PDX biopsies confirmed that CXCL12 was higher in tumor regions with fibroblast-clusters, while expression of CXCR4, the receptor of CXCL12 was higher in regions with KS signature clusters (Fig. 7G). This compartmentalized expression pattern of CXCL12 and CXCR4 indicates that the KS PDX system will be useful to further delineate the communication networks that promote proliferative expansion of KS infected endothelial cells in the tumor microenvironment.

## DISCUSSION

PDX models for other DNA virus-associated tumors, such as HPV-associated cervical cancer (*77*) and head and neck cancer (*78, 79*), HBV-associated hepatocellular carcinoma (*80, 81*), and EBV- associated NPC (*71, 82, 83*) have been previously reported. This is the first report establishing a KS PDX model for cutaneous KS. The KS PDX model supports the expansion of KSHV-infected spindle-shaped tumor cells for prolonged periods and maintain the cellular complexity of the tumor microenvironment, including KS, immune cells and other stromal cells that derive from the patient. The system is robust and the expansion of LANA+ cells was not found to be linked to clinical features or treatment history. Spatial transcriptomics is a powerful approach to query patient samples that are limited in size (*84*). This platform revealed that the PDX supported a shift towards more lytic gene expression beyond LANA, v-cyclin and Kaposin that was largely concurrent with a shift in a host gene expression profile conducive to angiogenesis. Last, one fibroblast-like lineage of cells isolated from the PDX were permissive to KSHV infection and produced a chemokine known to engage a chemokine receptor on infected endothelial cells.

KS is an atypical malignancy in that patient tumors do not maintain KSHV upon propagation in culture (*69, 85, 86*). We successfully maintained 27 P1 KS xenografts in immunodeficient mice from 103 d to 272 d until the recipient’s health deteriorated, requiring euthanasia. This mean 184 d timeframe is likely sufficient to examine potential interventions. The number of KSHV-infected endothelial cells increased in 14 out of 15 randomly examined P1 PDXs as compared to the original biopsy samples, accompanied by a robust detection of LANA expression in the Ki-67+ population cells. Interestingly, only a modest percentage of LANA+ cells were proliferative as determined by Ki-67 positivity in PDXs. The proliferative burst might have occurred in the earliest days after implantation, suggesting some type of homeostatic maintenance of infected cells in the PDX once the boundaries of the implant are reached at the later timepoints we examined. Regardless, what might be perceived as a failure of the skin KS PDX tumors to extend beyond the margins of the initial implant in the first or second passage in mice suggests that additional signals or tumorigenic events are required for the invasion of KS into surrounding tissue. We conclude that the skin KS PDX likely maintains the genetic stability reported for T1 stage KS (*87, 88*).

Multiple variables, including the clinical history of patients and implantation conditions, were evaluated for effects on the success rate of KS engraftment, based on the quantitative metric of an increase in LANA+ cell density from the input biopsy to the PDX explant. Recurrent KS previously treated with chemotherapy, antiretroviral therapy, immunomodulatory agents, and/or targeted therapies might be expected to show more aggressiveness than previously untreated KS. Patients with KICS have substantially elevated levels of cytokines, such as IL-6, and almost always have severe KS (*4*). However, the expansion of the LANA+ region in these different patient groups did not demonstrate statistically significant differences. One interpretation is that viral infection is a predominant driver in the expansion of infected areas. On evaluation of implantation factors, VEGF, a cytokine frequently detected in KS (*89*), is regarded as a key factor for shaping the pro-angiogenic KS tumor microenvironment (*90*). Given the trend towards a slight benefit in LANA+ expansion in VEGF-treated PDXs, we chose to retain VEGF supplementation moving forward.

The classification of KSHV latency-associated genes and lytic genes primarily stems from *in vitro* investigations with KSHV-infected PEL cell lines (*21, 91–93*). With specifically designed viral probes targeting five different KSHV transcripts, spatial transcriptomic analysis has revealed a more diverse expression pattern of KSHV genes in KS rather than the binary pattern observed in PEL cell lines (*34–36, 94–96*). Interestingly, the expression levels of all five transcripts studied here, including two lytic transcripts, were elevated in the PDXs, driving a shift from a predominantly latent infection in biopsy samples to a more lytic profile in PDXs. To some extent, this might reflect the flare of viral replication upon further loss of immune control in the PDX that recapitulates KS progression in patients with uncontrolled HIV and CD4^+^ T-cell lymphopenia (*97–99*). We did not detect viral DNA circulating in the blood, but future studies will evaluate if antiviral drugs block infected cell expansion in the PDX. If evidence for full productive infection occurs, the injection of uninfected cells may provide an opportunity for virus transmission and tumor growth, as recently reported in a xenograft model that included a mixture of circulating endothelial colony forming cells infected with KSHV and primary LEC (*50*).

Interestingly, we found multiple cell types interspersed in regions of infection in both biopsy and PDX tissues. To dissect which KSHV-infected cellular population in biopsy and PDX best represents the gene expression profile of KS, we identified a unique cluster with the expression of KS-signature genes and found that this cluster was separated from other cell types. The spots in this KS-signature cluster were mostly infected with KSHV. The KS cluster was most closely related to the gene expression profile of LECs and found proximal to the LECs that exhibited high viral gene expression in the PDX. This is consistent with the lymphatic origin proposed for KS spindle cells and, to some extent, supports the trajectory of viral reprogramming of lymphatic endothelial cells into KS cells (*8, 49, 63, 64, 100–102*).

Spatial transcriptomic analysis identified a number of DEGs between KSHV-uninfected and infected populations in biopsy and PDX samples, mainly from KSHV lytic clusters. This underscores KSHV lytic gene expression as a potential driver of reprogramming. The DEGs characterized in KS-signature and infected clusters were enriched in angiogenesis, extracellular matrix organization, endothelial cell development and differentiation, and cell growth, which align with the tumor biology of KS (*8, 103, 104*). Furthermore, the DEGs identified from spatial transcriptomic analysis of KS signature and KSHV-infected clusters also demonstrated selective downregulation of a pathway supporting lipid metabolism and transportation, though the biological relevance behind this observation remains to be further investigated. Notably, *MMP1*, *MMP12*, *MMP9*, and *CXCR4* were identified as the top DEGs associated with KSHV infection and the KS signature here, consistent with multiple datasets from tumors and infected endothelial cell infection systems (*36, 64, 76, 105*). These DEG expression levels were consistently upregulated in PDXs compared to matched biopsies, suggesting that the KS phenotype is amplified to some extent in PDXs. This further supports the utility of the KS PDX platform to test novel interventions.

Fibroblast cells were abundant in KS biopsies and PDX, characterized by gene profiles consistent with three functional subtypes of fibroblasts that had spatially distinct locations in the tumor. For instance, clusters predicted to be predominantly mesenchymal fibroblasts were more proximal to KS signature clusters compared to secretory fibroblasts. Two independent lineages of cells were derived from KS-PDX. These KSX-476 and KSX-488 derived cells exhibited variations in the level of PDGFR-α, PDGFR-β, SMA, FAP, and S100A4, markers shared between TAFs and mesenchymal stem cells (*72, 73, 106–109*). In both HIV-negative and HIV-positive KS tissues, PDGFR-α and SMA are co- expressed with LANA in KS spindle cells (*64*). TAFs are a critical component of the tumor microenvironment in multiple cancers including pancreatic and breast cancer. TAFs directly interact with tumor cells and shape the tumor milieu by secreting cytokines, chemokines, and enzymes that recruit immune cells and reorganize the extracellular matrix (*72, 110*). Future investigations are required to determine the role of different types of fibroblasts in KS.

KS-PDX cells were permissive for *de novo* infection but did not support full productive replication over an 8-day infection time course. KSHV infection differentially impacted host gene expression in KSX-476 and KSX-488; infection induced CXCL12 production from KSX-476. Engagement of CXCL12 with its receptors, CXCR4 and CXCR7, activates several signaling pathways, including PKC, MAPK, Src, and PI3K/AKT, leading to enhanced cell survival, migration, and proliferation (*75, 111*). CXCL12, CXCR4, and CXCR7 are elevated in KS compared to normal skin (*76*). KSHV infection of LECs induces CXCR4 (*64*) and mTOR, a downstream target in the CXCL12-CXCR4 signaling axis (*111*), is activated upon KSHV infection of LECs and KS tumor models *in vivo* (*51, 112–114*). The spatial transcriptomic analysis reported herein revealed a compartmentalized expression pattern of CXCL12 in fibroblast-rich regions and CXCR4 in tumor regions dominated by KS signature cells, that is more pronounced in the PDX. Taken together, the CXCL12-CXCR4 pathway orchestrated by multiple cells in the tumor microenvironment may promote KS progression and highlights the utility of the KS PDX for further study of cell-cell communications in KS.

Multiple animal models are reported as surrogate systems to study KS tumor biology and KSHV replication *in vivo*, and each has its strengths and limitations. KSHV vGPCR-immortalized human endothelial cells and vGPCR transgene expression in mice under the control of the vascular endothelial cell-specific promoter induced angio-proliferative tumors that share some features of human KS such as the occurrence spindle-shaped endothelial cells and infiltration of mouse immune cells (*30, 39, 115–117*). However, this single gene model does not allow for the investigation of viral gene functions within the context of a full viral infection. Xenograft models using KSHV-infected cells of rodent or human origin reproduced KS-like vascular tumors in immunodeficient mice, presenting with KSHV-infected endothelial spindle cells (*46–50*). Introduction of the KSHV genome into mouse bone marrow endothelial cells led to a KS-like gene expression pattern in spindle cell sarcomas, and RNAi targeting vGPCR reduced angiogenesis and tumor growth (*46*). Cell-line based models enable the investigation of gene-specific function or analyze the impact of small molecules on KS (*118, 119*). However, these approaches do not allow for the study of the pathological impact of different clinical strains of KSHV, nor do they recapitulate the complex cellular composition observed in KS tumors *in vivo*. The recent report of a single transgenic mouse line carrying the entire KSHV genome that develops angiosarcoma-like tumors with KSHV gene expression could be a valuable system to test virus-specific therapeutics (*51*). However, the integration of the KSHV genome in this model restricts the study of the biological significance of the fully productive phase of the KSHV lifecycle, and the genetic tractability of this system and transfer of tumor cells to naïve mice has not been reported. The skin KS PDX system enables the query of virus-driven processes and cell interactions in the primary cells of a patient tumor that will complement and extend existing models.

The skin KS-PDX model does have limitations. The restriction of tumor growth to the boundaries of the initial implant restricts passaging and larger cohort studies. No KSHV+ cell line was established from KS-PDX explants. We infer that the infected KS PDX cells are not truly transformed and retain dependence on the virus, and potentially the cellular environment, for persistence. The factors supporting viral-driven persistence of infected endothelial cells *in vivo* that are missing from cell culture models are not known. Last, the implantation times were prolonged but given the lack of a correlation between LANA+ cell expansion and time of implant, PDX of less than 100 days are expected to be successful. Regarding the spatial transcriptomic analysis, resolution was limited to ∼10 cells/spot. Single cell resolution is needed to link specific viral gene expression profiles to cell types and enable more accurate inferences of cell interactions. The limited sensitivity of low copy number transcripts likely leads to false negatives and an actual underestimation of viral and host reads. We expect newer technological advancements will overcome this limitation of the analysis. Despite these limitations, this is the first study, to our knowledge, that has implanted KS biopsies exclusively from PWH with varying clinical characteristics that has demonstrated persistence and expansion of the PDX.

Herein, we report the high fidelity of PDXs in supporting the expansion of KSHV-infected cells while also largely preserving the high dimensionality of the heterogeneous primary cells of KS tumors. The KS PDX provides a robust platform to test the role of infection and cellular factors in promoting expansion and tumor progression. Autologous transfer of patient PBMCs who have received KS-targeted immunotherapies is another potential pursuit. Future investigations will apply this model as an avatar model, wherein the mice with a KS-PDX will be treated in parallel with therapies used for the patient. We expect that this powerful approach will inform the mechanism of action that relates to the KS tumor response of the patient.

## MATERIALS AND METHODS

### Patient cohort and specimen collection

Individuals with KS treated in the National Institutes of Health Clinical Center under the care of the HIV and AIDS Malignancy Branch at the National Cancer Institute were included. Clinical and HIV characteristics were obtained at the time of biopsy collection. KSHV viral load (VL) per million peripheral blood mononuclear cells (PBMCs) was measured in DNA extracted from PBMCs by quantitative real-time polymerase chain reaction using K6 primers, as previously described (*120*).

Participants had a 6 mm punch biopsy of cutaneous KS. All participants were consented to protocols for tissue procurement (NCT00006518) and/or sequencing of KS and other KSHV- associated diseases (NCT03300830) to permit RNA sequencing of KS lesions for this study. Both protocols were approved by the NIH Institutional Review Board. All enrolled participants gave written informed consent in accordance with the Declaration of Helsinki.

### Experimental animals

NCI-Frederick is accredited by AAALAC International and follows the Public Health Service Policy for the Care and Use of Laboratory Animals. Animal care was provided in accordance with the procedures outlined in the “Guide for Care and Use of Laboratory Animals (National Research Council; 1996; National Academy Press; Washington, D.C.).” All study protocols were approved by the NCI at Frederick Animal Care and Use committee (Frederick, MD). NSG mice were bred at the NCI-Frederick animal facilities. HIL-6 NOG mice were obtained from Taconic (Cat. #13686, Germantown, NY).

### Processing and implantation of KS biopsy

NSG or hIL6 NOG mice were implanted subcutaneously with fresh KS skin tumor biopsy fragments from the clinical study (IRB #01C0038). A small fragment of the biopsy was excised and placed in 10% neutral buffered formalin for histology analysis. The remaining biopsy tissue was rinsed with 1X PBS biopsy fragment in a petri dish and (if large enough) dissected into fragments of ∼5 mm X 5 mm. Fragments were implanted subcutaneously in the mouse flank in a 1:1 ratio of serum-free RPMI medium:extracellular matrix (Matrigel solution, cat# 356234, Corning, Tewksbury, MA) containing 0.5 µg/ml human VEGF 165 (cat# 100-20, Peprotech, Cranbury, NJ). Starting on day 3 post-implant, mice were treated with rituximab at 25 mg/kg IP, 3 times per week in the first week, followed by once per week for 3 weeks. Mice were monitored daily, with caliper measurements and body weights recorded bi-weekly. Endpoints included weight loss greater than 20% of initial body weight, hunching, lack of movement, or any clinical signs of distress.

### Histology and Immunohistochemistry

Biopsy fragments and PDX tumors were fixed in 10% neutral buffered formalin for 48-72 hrs, embedded in paraffin, and 5 μm sections were transferred to glass slides for routine hematoxylin and eosin staining and immunohistochemistry (IHC). Primary antibodies used for staining included the following: HHV8/LANA (cat# PA0050, Leica, Bannockburn, IL), CD31 (1:200, cat# ab32457, Abcam, Cambridge, MA), CD34 (1:200, cat# MBS684197, MyBiosource, San Diego, CA), NUMA1 (1:75, cat# LS-B11047, Lifespan Biosystems, Seattle, WA), Ki-67 (1:250, cat# 9027, Cell Signaling Technology, Danvers, MA) and vIL-6 (1:3000, rabbit monoclonal antibody, Epitomics, Burlingame, CA (*121*). All IHC staining was performed on the Bond-Max autostainer (Leica). Sections were subjected to heat-induced epitope retrieval with citrate (Bond Epitope Retrieval Solution 1, Leica Biosystems; CD31, Ki-67, NUMA1) or EDTA (Bond Epitope Retrieval Solution 2, Leica; CD34, LANA) for 20 min prior to immunostaining. Slides were incubated with primary antibody for 15 min (LANA) and 30 min (CD31, CD34, Ki-67, NUMA1). Signal was detected using a Bond Polymer Refine Detection DAB (BPRD) kit (DS9800, Leica) with hematoxylin counterstaining. Brightfield slides were scanned at 20x using an Aperio AT2 scanner (Leica).

### Immunofluorescence

Dual immunofluorescent staining for LANA and Ki-67 was performed on paraffin-embedded tumors. After 20 minutes of HIER with EDTA (Bond Epitope Retrieval Solution 2, Leica), tissue was blocked with Ultracruz (sc-516214, Santa Cruz Biotechnology, Santa Cruz, CA) for 60 min and the antibodies were incubated at room temperature for 15 min (LANA) and 60 min (Ki-67). Slides were incubated with a cocktail of Alexa fluor-conjugated secondary antibodies for 30 min (1:250, cat# 4409 and 4419, Cell Signaling Technology).

Triplex immunofluorescent staining was performed at the Molecular Histopathology Laboratory (Frederick National Laboratory for Cancer Research) for CD34, LANA, and IBA1 using the Bond- RX/3 autostainer (Leica) with the BPRD kit (DS9800, Leica) reagents. After 15 min of HIER EDTA (Bond Epitope Retrieval Solution 1, Leica Biosystems), peroxide block was applied with a mouse Ig block and normal horse serum (cat# S-2000, Vector, Newark, CA), CD34 antibody was applied at ambient temperature for 30 min (1:50, cat# NCL-L-END, Leica, Bannockburn, IL) followed by an anti-mouse secondary antibody conjugated to HRP and visualized with Opal 690 (cat#OP-001006, Akoya, Marlborough, MA). CD34 was quenched using the peroxide block. LANA (1:800, cat#MABE1109, Millipore Sigma, Darmstadt, Germany) was incubated at ambient temperature for 30 min and detected with HRP-conjugated goat anti-rat and visualized with Opal 620 (cat#OP-001004, Akoya, Marlborough, MA). The LANA signal was quenched using the Leica kit’s peroxide block. The BPRD kit’s polymer reagent was used to detect IBA1 (1:500, cat#CP290, Biocare, Pacheco, CA), which was applied at ambient temperature for a 30 min incubation and visualized with Opal 520 (cat#OP-001001).

ProLong Diamond Antifade Mountant with DAPI (cat# P36971, Invitrogen, Carlsbad, CA) was applied to all fluorescent slides (aqueous mount) prior to the coverslip. Immunofluorescent slides were scanned at 20x using a PhenoImager HT (Akoya Biosciences, Marlborough, MA) at the Molecular Histopathology Laboratory (Frederick National Laboratory for Cancer Research). Quantitative image analysis was performed on manually annotated tumor regions using the HALO image analysis platform’s Cytonuclear v.2.05 and CytoNuclear FL v2.0.12 algorithms (Indica Labs, Corrales, NM).

### Capturing spatial transcriptomics of KS and PDX

Visium Gene Expression slides (cat# 1000338, 10X Genomics, Pleasanton, Ca) with investigator placed sections from FFPE treated samples were deparaffinized, H&E stained, imaged and decrosslinked. The tissue was permeabilized and 10X Genomics human transcriptome V1 probes (10X Genomics) were hybridized to the mRNA overnight. The probes were ligated, released from the tissue, captured on the spatially barcoded spots and subjected to extension followed by denaturation per manufacturer protocol. The full-length probes were subjected to real-time PCR quantification to determine optimal sample index PCR cycles to complete the Illumina-ready library. Final library quality was assessed using the LabChip GXII HT (PerkinElmer, Wellesley, MA) and libraries were quantified by Qubit (Thermo Fisher, Pleasanton, CA). Pooled libraries were subjected to paired-end sequencing according to the manufacturer’s protocol (Illumina NovaSeq 6000). Bcl2fastq2 Conversion Software (Illumina, San Diego, CA) was used to generate de-multiplexed FASTQ files. These procedures were performed by the University of Michigan Advanced Genomics Core.

### Aligning FASTQ files after removing mouse and ambiguous reads from spatial transcriptomic database

The original FASTQ files were generated through 10X Visium pipeline with the respective TIFF image file using v1 probe with 5 KSHV spiked in the probes. To remove the ambiguous and mouse reads, previous methods were adapted to the spatial datasets that are embedded with additional barcode sequences in the reads. The preprocessing part of the Xenomake pipeline (*122*) was applied, which removed the spatial barcode/UMI tag and trims adapter/polyA from the reads. Thereafter, Xenome was applied to determine if the source of the read is the graft (i.e., human) or the host (i.e., mouse). Ambiguous or mouse reads from the paired FASTQ files were removed based on Xenome calls, keeping only the reads whose source was the human xenograft or the virus. Processed FASTQ files were then aligned to a custom reference with KSHV genome (NC_009333.1) added using SpaceRanger “count” module (version 3.0.0).

### Removal of Low Read Count Cluster and Batch Correction

Low read spots were removed using Seurat (version: Seurat_5.1.0) in R. Samples were merged into a single Seurat object using a standard pipeline (by normalizing, finding top 2000 genes, scaling data, RunPCA, RunUMAP). Batch effects were removed using the Harmony package (version: harmony_1.2.0) and then analyzed using Seurat’s standard pipeline (RunUMAP, FindNeighbors, choosing the top 30 dimensions). Seurat FindCluster (default parameter notably resolution=0.8) was utilized to generate 13 clusters. The cluster with a significantly lower median UMI count of 239 from the other clusters was removed. KSHV gene expression was analyzed using raw UMI counts plus one by log2 transformation.

### Cell Decomposition and Analysis

The cell composition was analyzed using the CARD package (CARD_1.1) (*62*). Five normal samples (GSE130973) were provided as reference. The original read counts (not normalized) were passed to the CARD algorithm. CARD assigned portions for each spot to 11 cell types plus an “unknown” cell type category. By adding the portions of some cell types, categories including Immune cells: macrophage/DC, T cells, and plasma cells; Endothelial cells, lymphatic and vascular endothelial cells; Keratinocytes, differentiated and undifferentiated keratinocytes; stromal cells, pericytes, and melanocytes were generated. Fibroblasts, erythrocytes, and “Unknown” remained as their own specific categories. Data was visualized using heatmap and geom_bar in the ggplot R package (ggplot2_3.5.1).

### Clustering of hierarchical integrated datasets

Data processing and visualization was performed in R using Seurat (version 5.1.1.0), scType (*123*), CellChat (version 1.6.1) (*124*), ComplexHeatmap (version 2.15.4), scCustomize (version 2.1.2) (doi 10.5281/zenodo.5706430), clusterProfiler and enrichplot (version 1.18.4) (version 4.13.0, <u>10.1089/omi.2011.0118</u>, <u>10.1016/j.xinn.2021.100141</u>.).

Normalization and variance stabilization was performed using the SCTransform function. Pairwise hierarchical integration was performed in two steps using the SelectIntegrationFeatures, FindIntegrationAnchors functions Integrate Data. First, biopsies and PDX samples were integrated, subsequently integration of combined biopsy and PDX objects was performed. After gene normalization via ScaleData, dimensionality reduction was performed through PCA. Clustering was performed at a resolution of 1.1 using FindNeighbors and FindClusters and visualized using tSNE and UMAP, resulting in 16 clusters with 10 clusters present in all samples.

### Cell type annotation of spatial transcriptomics datasets

Predictions of dominant cell type signatures per cluster were performed using scType with cell-specific marker genes for 14 cell types present in skin (*61*) and 3 additional cell types manually annotated based on established marker genes (B cells, Plasma Cells) and a *de novo* set of five marker genes, ADAMTS4, ADAM19, FLT4, STC1, CLEC4M, for KS tumors (KS Signature), . The following process was used to identify the genes for the KS Signature. Thirty-three DEGs from previous studies (*35, 36*) were considered candidates for the KS Signature. Using these genes, a logistic regression model (glm function stats R package) was trained to distinguish between KSHV+ and KSHV- spots for each matched pair of Biopsy and PDX. The KS Signature was the intersection of the genes found as significant (p<0.001) by both models. To validate the method for generating KS Signature, each model was re-trained using the significant genes identified in its training and evaluated on the unseen matched pair by the model. The Area Under the Curve (AUC) of the Receiver Operating characteristic of both models trained only on significant genes was around 77%, and indicates the signature’s stability. All downstream analyses were performed excluding keratinocytes to avoid bias based on dominant keratinocyte markers.

### Differential gene expression analysis of spatial transcriptomics datasets

Differentially regulated genes between populations based on cell types were predicted using the FindAllMarkers function, restricted to genes detected in a minimum of 10% in either population. The 25 top up- and downregulated markers for the KS signature cluster with adjusted p-values <0.001 and expression in a minimum of 50% cells in the KS signature population (for up-regulated genes) or 50% cells in the comparison group (for down-regulated genes) were used for visualization with the scCustomize Clustered_DotPlot function. Gene enrichment analysis and visualization was performed using the clusterProfiler and enrichplot R packages for up- or downregulated genes (cutoff 2-fold change, adjusted p-value <0.001) in the KS signature cluster. Linkages of genes and biological processes were visualized using cnetplot. Differentially regulated genes between populations based on infection status were predicted using the FindMarker function using comparisons with uninfected spots, restricted to genes detected in a minimum of 10% in either population. The 15 top up- and downregulated markers for each comparison (latent, lytic, mixed vs uninfected) with adjusted p-values <0.001 were used for visualization with the scCustomize Clustered_DotPlot function. Gene enrichment analysis using clusterProfiler was performed for up- or downregulated genes found in at least one comparison (cutoff 2-fold change, adjusted p-value <0.001). Linkages of genes and biological processes were visualized using cnetplot.

### Comparison of differentially regulated genes with published bulk RNA sequencing datasets

Raw count matrices for datasets GSE147704 (*35*), GSE241095 (*36*) were downloaded from the Gene Expression Omnibus database. Normalization and differential expression analysis was performed in the NIH Integrated Analysis Portal (NIDAP) using R programs developed on Foundry (Palantir Technologies). Raw counts were filtered to retain genes that had at least three samples with non-zero CPM counts in at least one group. Differential expression of genes of log2- CPM transformed raw counts was performed using the Limma Voom R package. The overlap of differentially regulated genes (cutoff 2-fold change, adjusted p-value <0.001) between different data sets was analyzed and visualized using InteractiVenn (*125*) and the ComplexHeatmap package (version 2.15.4) (10.1093/bioinformatics/btw313, 10.1002/imt2.43).

### Isolation, culture, and infection of <u>KS</u> PD<u>X</u> derived cells (KSX)

Matrigel solution (5 mg/ml) was added to each well of a 12-well plate. PDX explant was trimmed to 2 mm^2^ and seated on Matrigel layer and submerged with 500 µl of Matrigel solution. After 30 min at 37°C, embedded PDX explant was overlaid with 2 ml of KSX culture medium (EGM-2 supplemented with 10 µM Y-27632 [cat# ALX-270-333, Enzo, Farmingdale, NY]). 1 ml of fresh KSX medium was replaced every 2 d until cells migrated out of the PDX tissue and formed a confluent cell layer. After washing with 500 µl of prewarmed 1X PBS, the KS PDX cell layer was disrupted and dissociated by treating with 500 µl of Dispase solution (cat# 354235, Corning) diluted in 1X PBS at 1.5 U/ml for 30-45 min at 37°C. KS PDX cells were subsequently separated from the remaining PDX tissue by passing through a 70 µm strainer (cat# 431751, Corning), pelleted, and seeded in 2 wells without Matrigel on a 12-well plate for continued expansion. KSX- 476 and KSX-488 were maintained in KSX culture medium and passaged by treating with Accutase Cell Dissociation Reagent (cat# A1110501, Thermo Fisher) after washing with 1X PBS. KSX-476 and KSX-488 were seeded at 1.5 X10^5^ cells per well with EGM-2 medium in a 24-well plate 18 hr prior to spinoculation with KSHV BAC16 (*126*) MOI=1 at 1800 rpm for 45 minutes at 37°C, and overlaid with 1 ml of EGM-2.

### RNA extraction and quantitative reverse transcriptase PCR

Total RNA from mock and KSHV infected KSX-476 and KSX-488 cells was extracted with RNeasy Plus kit (Qiagen) followed by DNase digestion (cat# EN0525, Thermo Fisher). DNase treated RNA was reverse transcribed to cDNA using the SuperScript IV First-Strand Synthesis system (cat# 18091050, Invitrogen) following manufacturer instructions. Relative transcript levels of selected cellular genes were determined by quantitative real-time PCR with gene-specific primers and PowerUp SYBR Green Master Mix (Applied Biosciences) in a QuantStudio 3 Real-Time PCR System (Applied Biosciences). Sequences of primers used are as follows: forward primer –GCTCGAATCCAACGGATTTG and reverse primer AATAGCGTGCCCCAGTTGC for *ORF26*; forward primer– CCCTGAGCCAGTTTGTCATT and reverse primer – ATGGGTTAAAGGGGATGATG for *ORF50*; forward primer TGGACATTATGAAGGGCATCCTA and reverse primer –CGGGTTCGGACAATTGCT for *ORF57*; forward primer –TTAGAAGTGGAAGGTGTGCC and reverse primer TCCTGGAGTCCGGTATAGAATC for *ORF59*; forward primer – TGGTCGGCGGTTCAGTCATCAA and reverse primer –GCGGCCGCTAAGAAAATCGA for *K8.1*; forward primer –GGATGCCCTAATGTCAATGC and reverse primer – GGCGATAGTGTTGGGAGTGT for *ORF71*; forward primer – GCTGATAATAGAGGCGGGCAATGAG and reverse primer – GTTGGCGTGGCGAACAGAGGCAGTC for *ORF72*; forward primer – GCAGACACTGAAACGCTGAA and reverse primer –AGGTGAGCCACCAGGACTTA for *ORF73*; forward primer –ATTCTCAACACTCCAAACTGTGC and reverse primer – ACTTTAGCTTCGGGTCAATGC for *CXCL12;*forward primer – TTGATTTTGGAGGGATCTCG and reverse primer –GAGTCAACGGATTTGGTCGT for *GAPDH*.

### Immunoblot

Total cell extracts in Tris-Glycine SDS Sample Buffer (cat# LC2676, Thermo Fisher) were electrophoresed in 4–20% SDS-polyacrylamide gels (cat# XP04200BOX, Thermo Fisher), transferred to nitrocellulose membranes, and blotted with indicated antibodies including mouse anti-CD31 (cat# 3528; Cell Signaling Technology), rabbit anti-PDGF Receptor α (3174; Cell Signaling Technology), rabbit anti-PDGF Receptor β (cat# 3169; Cell Signaling Technology), rabbit anti-α-Smooth Muscle Actin (cat# 19245; Cell Signaling Technology), rabbit anti-VEGF Receptor 3 (cat# 2638; Cell Signaling Technology), rabbit anti-FAP (cat# 66562, Cell Signaling Technology), rabbit anti-S100A4 (cat# 13018, Cell Signaling Technology) and rabbit anti-GAPDH (cat# 2118; Cell Signaling Technology). Signals were detected using enhanced chemiluminescence and acquired on iBright Imaging Systems (Thermo Fisher).

### Enzyme Linked Immunosorbent Assay (ELISA)

The ELISA quantitating extra-cellular CXCL12 level was performed with Human CXCL12/SDF- 1 DuoSet ELISA kit (cat# DY350-05, Bio-Techne, Flowery Branch, GA) following manufacturer instructions. Briefly, 100 μL of conditioned medium from mock or KSHV infected KSXs along with serially diluted standards were loaded on the plate coated with capture antibody against CXCL12. After incubating for 2 h at room temperature, the plate was washed, incubated with CXCL12 detection antibody for 2 h and Streptavidin-HRP for 20 min. Substrate was add in each well for incubation for 20 min before reaction was stopped. The plate was analyzed for optical density at 450 nm wavelength (Gen5, BioTek, Winooski, VT).

## Statistical analysis

All data was analyzed using GraphPad Prism software (GraphPad Software version10, http://www.graphpad.com<u>, La Jolla, CA</u>). Paired analysis, followed by two-tailed t test was applied for analysis of changes between the biopsy and PDX. Unpaired, two-tailed t test was applied for comparisons between two independent groups. Simple linear regression analysis was applied to examine correlations between variables. Gene expression was evaluated as unpaired, two-tailed t test on log2 normalized counts or two-way ANOVA followed by Dunnett’s multiple comparisons. P values are defined in figure legends.

## List of Supplementary Materials

### Materials and Methods

Supplementary Fig. 1. Passage 2 PDX explants do not expand but retain features of KS.

Supplementary Fig. 2. Cell-type identification and viral gene expression in clusters.

Supplementary Fig. 3. Gene set enrichment analysis and pathways of DEGs.

Supplementary Fig. 4. KSHV lytic reactivation is impaired upon inhibition of Rho kinases.

Supplementary Fig. 5. KSX cells are human tumor associated fibroblast-like cells that are KSHV negative but highly permissive to viral entry.

Supplementary Fig. 6. Spatial distribution of CXCL12 (upper) and CXCR4 (lower) SCT normalized expression levels in fibroblast and KS signature clusters in biopsy/PDX pair 2.

Supplementary Table 1. Distribution of KSHV transcript and infection status in biopsy and PDX.

Supplementary Table 2. Gene signatures to define cell clusters.

Supplementary Table 3. Short Tandem Repeat (STR) profiles.

## Acknowledgments

We thank Rajaa El Meskini, Vickie Marshall, and Denise Whitby for technical advice and discussions.

## Author contributions

Conceptualization: XL, ZWO, LTK Methodology: XL, AD, LB, AG, BA, TT, WC, JZ

Investigation: XL, ZWO, AD, LB, AG, BA, WC, RM Visualization: XL, ZWO, LB, AG, BA, WC, LTK Funding acquisition: RY, JZ, RR, LTK

Project administration: XL, ZWO, LTK Supervision: ZWO, RY, JZ, RR, LTK

Writing – original draft: XL, ZWO, LB, AG, BA, RR, LTK

Writing – review & editing: XL, ZWO, LB, AG, BA, WC, KL RR, LTK

## Data and materials availability

Data will be made publicly accessible following publication.

