## Supplemental Files for "Mapping herpesvirus-driven impacts on the cellular milieu and transcriptional profile of Kaposi sarcoma in patient-derived mouse models"

**Supplementary Materials:**

**Short tandem repeat analysis**

Cells for STR analysis were prepared with FTA Sample Collection Kit and submitted to ATCC cell authentication testing service. Briefly, KSX were detached from adherence with Accutase. The single cell solution of KSXs along with PBMC from matching donor were counted, pelleted and resuspended in 1X PBS at cell density of $1 \times 10^6$ cells/ml. 40 µl of cell suspension was spotted on the sample collection card and air-dried before being packed. Seventeen short tandem repeat (STR) loci plus Amelogenin, the gender determining locus, were amplified using PowerPlex® 18D Kit from Promega. The cell sample was processed on ABI Prism® 3500xl Genetic Analyzer. Data were analyzed using GeneMapper® ID-X v1.2 software (Applied Biosystems, College Station, TX). Appropriate positive and negative controls were included and confirmed for each submitted sample.

**Western Blot**

Cell lysates were electrophoresed in 4–20% SDS-polyacrylamide gels, transferred to nitrocellulose membranes, and blotted against KSHV and cellular proteins with primary antibodies including mouse anti-ORF45 (cat# sc-53883; Santa Cruz), mouse anti-K8.1 (cat# sc-65445; Santa Cruz) and rabbit anti-GAPDH (cat# 2118; Cell Signaling Technology). Signals were detected using enhanced chemiluminescence and acquired on iBright Imaging Systems.

**Immunofluorescence**

Cells were fixed at room temperature for 15 min with fixation solution (cat# 554722, BD Bioscience), washed with 1X Perm/Wash buffer (554723, BD Bioscience) and incubated with NUMA1 antibody (cat# LS-B11047, LSBio) for 1 h at room temperature. After washing, cells were further incubated with Alexa Fluor 488 conjugate anti-rabbit antibody (cat# A-11008, Thermo Fisher) for 45 min and then mounted using Prolong Gold Antifade with DAPI (4′,6-diamidino-2-phenylindole) (cat# P36935, Thermo Fisher Scientific) for imaging under microscopy. The pictures were taken from representative fields for each cell type.

**DNA extraction and qPCR**

Total cellular DNA from iSLK, iSLK-RGB, KSX-476 and KSX-488 cells was isolated with DNeasy Blood & Tissue Kits (Qiagen) following the manufacture's manual. KSHV DNA was quantitated using quantitative-PCR by amplifying KSHV *K9* gene with forward primer, 5′-GTCTCTGCGCCATTCAAAAC-3′, and reverse primer, 5′-CCGGACACGACAACTAAGAA-3′.

**Quantitative PCR for angiogenesis genes**

Taqman PCR was performed on TaqMan™ Array Human Angiogenesis plate (cat# 4414071, Thermo Fisher) with lyophilized primers and probe targeting selected genes involved in human angiogenesis or housekeeping gene. For qRT-PCR, 10 µl of Taqman Fast Advanced Master Mix (cat# 4444557, Applied Biosystems) was mixed with 10 µl of cDNA in each well to reconstitute the lyophilized primers and probe. The cDNA used for each plate was a mixture of equal amount of cDNA obtained from the three biological repeats described in Fig. 7 D and E. qRT-PCR was run on QuantStudio 3 Real-Time PCR System (Applied Biosciences), data were captured and presented in the form of Ct (cycle threshold) value.

**Supplementary Table 1**

Distribution of KSHV transcript and infection status in biopsy and PDX

| | **Biopsy 1** | **PDX 1** | **Biopsy 2** | **PDX 2** |
|---|---|---|---|---|
| infected of all spots | 30.20% | 99.48% | 27.16% | 95.03% |
| *ORF72* of all spots | 5.87% | 37.61% | 3.40% | 47.42% |
| *K12* of all spots | 13.18% | 59.94% | 8.42% | 74.81% |
| *ORF75* of all spots | 1.91% | 28.09% | 2.16% | 33.67% |
| *K8.1* of all spots | 0.53% | 25.19% | 0.12% | 17.68% |
| *PAN* of all spots | 16.15% | 97.36% | 15.73% | 87.17% |
| Latent of infected | 43.39% | 0.22% | 31.30% | 4.08% |
| Lytic of infected | 1.39% | 28.43% | 0.17% | 19.94% |
| Mixed of infected | 55.22% | 71.35% | 68.53% | 75.98% |

**Supplementary Table 2.**

Gene signatures to define cell clusters.

| **Cell types** | **Marker genes** |
|---|---|
| Keratinocytes (undiff) | *KRT5, KRT14, TP63, ITGB1, ITGA6* |
| Keratinocytes (differentiated) | *KRT1, KRT10, SBSN, KRTDAP* |
| Fibroblasts | *PDGFRA, LUM, DCN, VIM, COL1A2* |
| Fibroblasts (secretory-reticular) | *WISP2, SLP1, CTHRC1, MFAP5, TSPAN8* |
| Fibroblasts (proInflammatory) | *CCL19, APOE, CXCL2, CXCL3, EFEMP1* |
| Fibroblasts (secretory-papillary) | *APCDD1, ID1, WIF1, COL18A1, PTGDS2* |
| Fibroblasts (mesenchymal) | *ASPN, POSTN, GPC3, TNN, SFRP1* |
| Melanocytes | *PMEL, MLANA, TYRP1, DCT* |
| Macrophages/ DC | *AIF1, LYZ, HLA-DRA, CD68, ITGAX* |
| VEC | *SELE, CLDN5, VWF, CDH5* |
| LEC | *PROX1, CLDN5, LYVE1* |
| Pericytes | *ACTA2, RGS5, PDGFRB* |
| Erythrocytes | *HBA1, HBA2, HBB* |
| T cells | *CD3D, CD3G, CD3E, LCK* |
| B cells | *CD19, FCGR3A, CD38, CD24, MS4A1* |
| Plasma cell | *CD27, CD38, SDC1, XBP1, PRDM1* |
| KSHV+ | *K8-1, T1-1, ORF72, ORF75, HHV8GK18-gp79* |
| KS signature | *ADAMTS4, ADAM19, FLT4, STC1, CLEC4M* |

**Supplementary Table 3.**

| | **Sample Type** | | | | | |
|---|---|---|---|---|---|---|
| ***Locus**** | **TB133 patient PBMC** | | **KSX-476 PDX cell culture 1** | | **KSX-488 PDX cell culture 2** | |
| *TH01* | 6 | 7 | 6 | 7 | 6 | 7 |
| *D5S818* | 11 | 11 | 11 | 11 | 11 | 11 |
| *D13S317* | 10 | 11 | 10 | 11 | 10 | 11 |
| *D7S820* | 12 | 12 | 12 | 12 | 12 | 12 |
| *D16S539* | 11 | 11 | 11 | 11 | 11 | 11 |
| *CSF1PO* | 12 | 13 | 12 | 13 | 12 | 13 |
| *Amelogenin* | X | Y | X | Y | X | Y |
| *vWA* | 16 | 16 | 16 | 16 | 16 | 16 |
| *TPOX* | 8 | 12 | 8 | 12 | 8 | 12 |

* STR in loci of *D3S1358, D21S11, D18S51, Penta_E, Penta_D, D8S1179, FGA, D19S433, D2S1338* were tested with matching profile between samples but not disclosed herein.

**Supplementary Fig. 1**

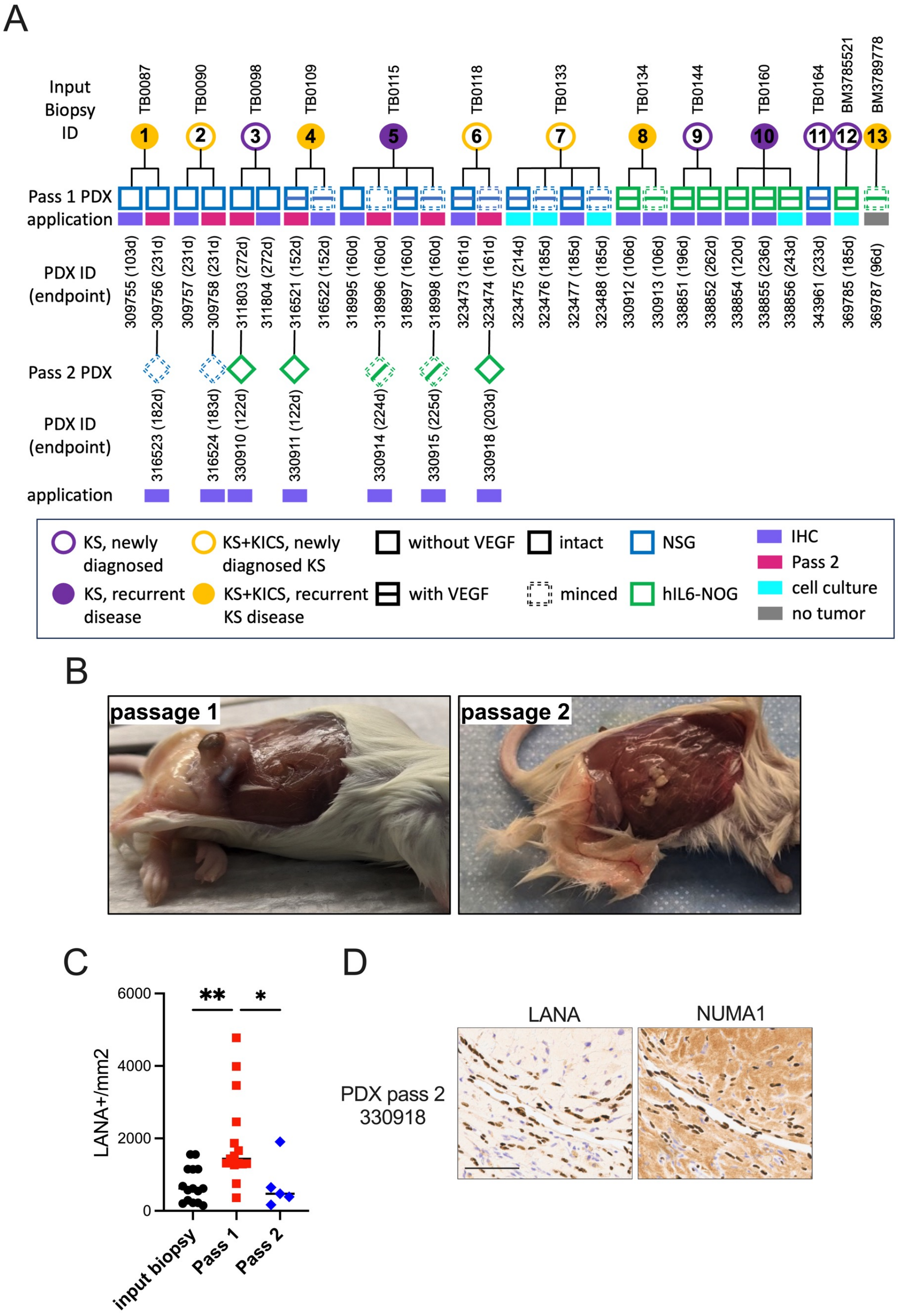

**Supplementary Fig. 1. Passage 2 PDX explants do not expand but retain features of KS.**

(**A**) As described in Fig 3A, here incorporating passage 2 (P2) PDX, a summary of all the KS biopsy (circles) and matched PDX explants (squares) described in this study, de-identified code for biopsy tissue (above circles) and recipient mouse ID (below squares) with indicated implantation times in parentheses. Clinical features of the patients (presentation of KS alone or KS in combination with KS inflammatory cytokine syndrome (KICS), primary or recurrent KS tumors) are indicated in the legend. Implantation variables included extracellular matrix supplemented without or with vascular endothelial growth factor (VEGF), leaving the tumor intact or minced, and implantation in NSG mice or human IL-6 transgenic NOG mice, as indicated in legend. **(B)** P1 PDX # 309755 (left panel; implant from biopsy TB0087), and corresponding P2 PDX #316523 (right panel; implant from minced P1 PDX). **(C)** Cells positive for LANA by IHC per tissue area ($mm^2$) were quantified in the biopsy and the corresponding P1 and P2 PDX explants; **, $p<0.01$; *, $p<0.05$, one-way ANOVA followed by Dunnett's multiple comparisons. **(D)** Two P2 PDX tumors were analyzed for proximity of LANA+, NUMA1+ cells to vessel (top panels) and vascular-like channels (bottom panels); scale bars = 100 μm.

**Supplementary Fig. 2**

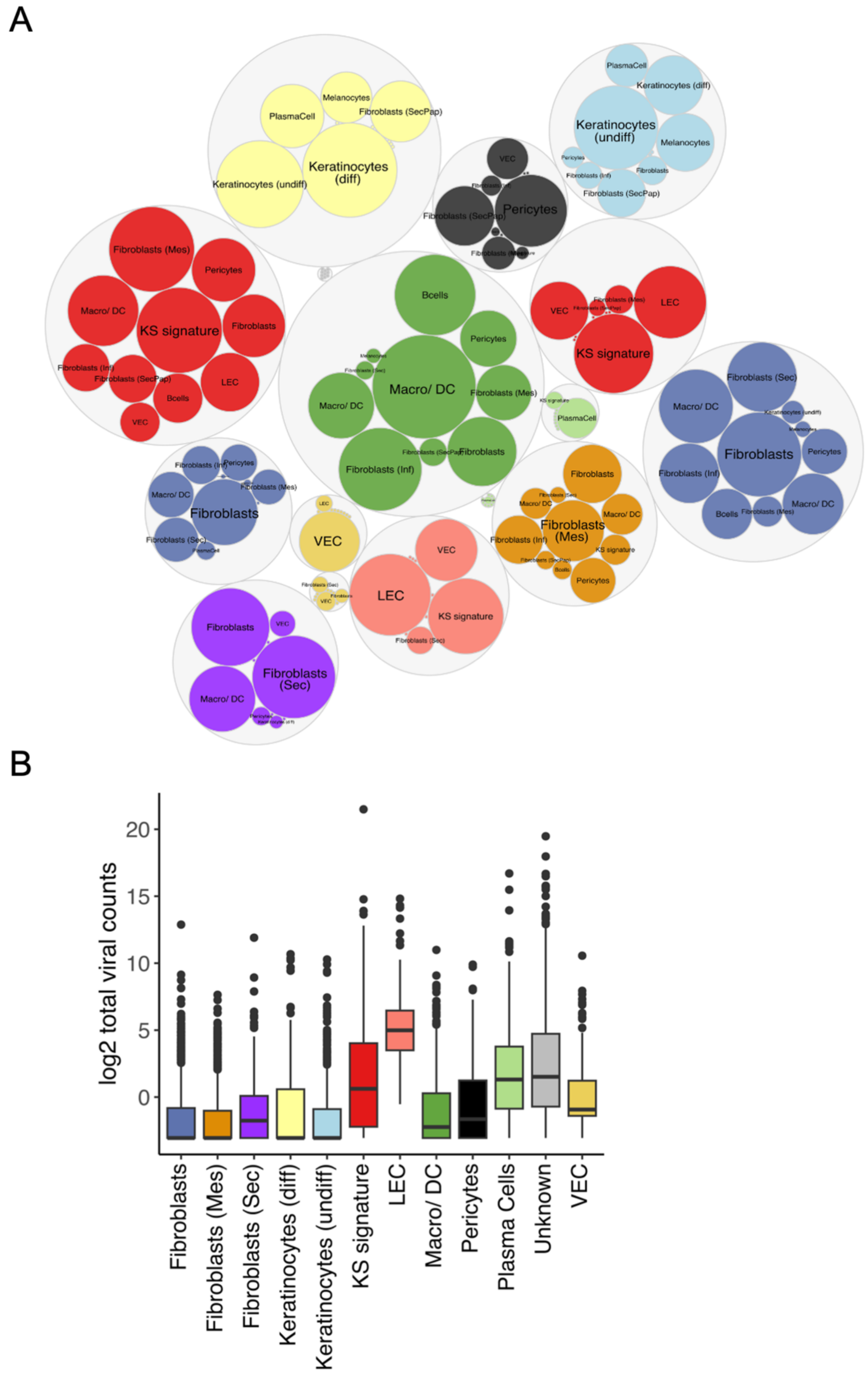

**Supplementary Fig. 2. Cell-type identification and viral gene expression in clusters.**

**(A)** Automated cell type identification based on marker genes for 12 cell types was performed using scType and scType scores for all cell types considered by scType for cluster annotation are shown in a bubble plot. Grey, outer bubbles are proportional to cluster size, inner bubbles signify scType scores for each cell type. The largest bubble and colors of bubbles in each cluster correspond with the assigned cell type. (**B**) $Log_2$ expression levels of total KSHV transcripts are presented as box plots for each cell type.

### Supplementary Fig. 3

A. Pathway and network analysis of upregulated DEGS in KS signature cluster

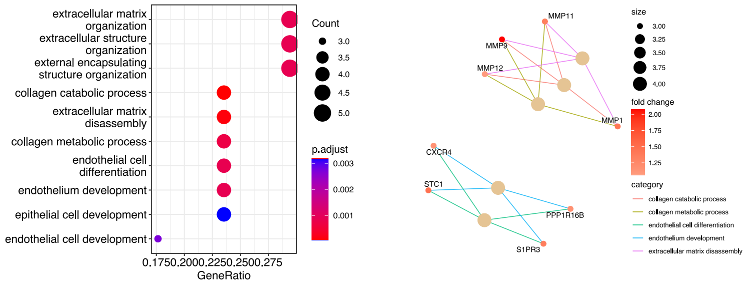


B. Pathway and network analysis of downregulated DEGS in KS signature cluster

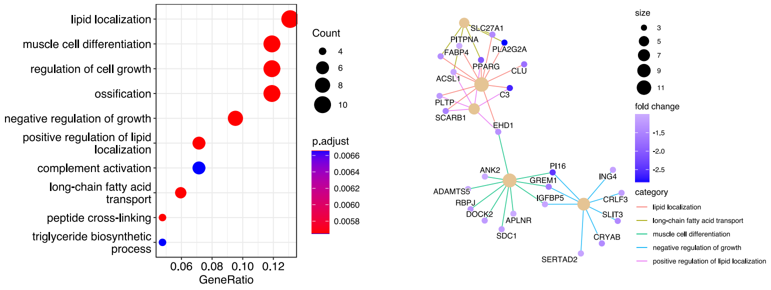


C. Pathway and network analysis of upregulated DEGS in KSHV infection

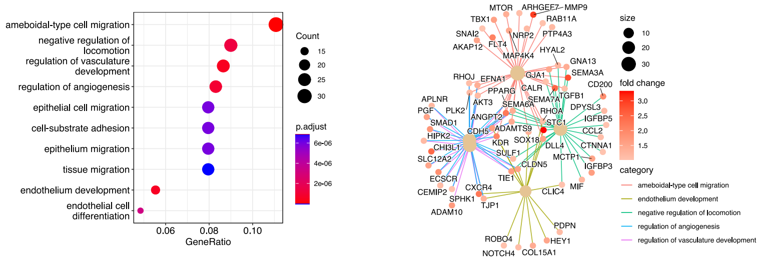


D. Pathway and network analysis of downregulated DEGS in KSHV infection

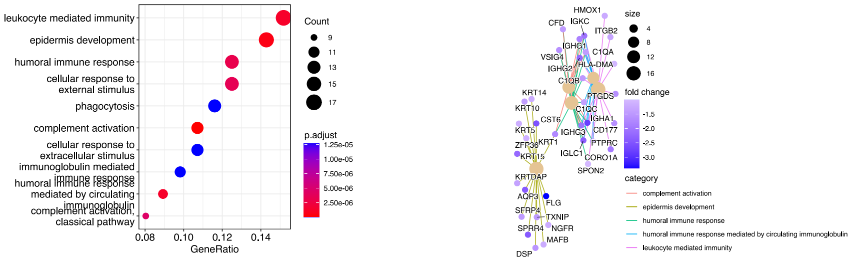


E.

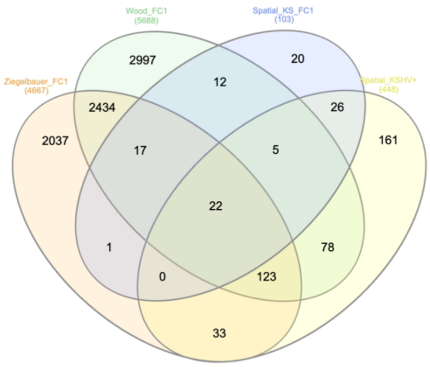

**Supplementary Fig. 3. Gene set enrichment analysis and pathways of DEGs.**

**(A-B)** Gene set enrichment analysis using the gene ontology (GO) aspect biological process of differentially expressed genes (DEGs) upregulated (**A**) or downregulated (**B**) in the KS signature cluster compared to other clusters at a cutoff of 2-fold change, adjusted p-value <0.001 (left panel). Colors depict enrichment scores (*e.g.* p values), gene counts, the number of genes enriched within the specified GO terms, are indicated by circle size and gene ratios, the ratio of DEGs to total genes within the specified GO terms, are displayed on the x-axis. Gene-concept networks of top 5 enriched GO terms. Colored connections depict GO terms, nodes sizes indicate gene counts within the specified GO terms and fold changes in the KS signature cluster compared to the other clusters are indicated by color. (**C-D)** Gene set enrichment analysis using the GO aspect biological process of DEGs upregulated (**C**) or downregulated (**D**) in infected spots compared to uninfected spots at a cutoff of 2-fold change, adjusted p-value <0.001 (left panel). Colors depict enrichment scores (*e.g.* p values), gene counts, the number of genes enriched within the specified GO terms, are indicated by circle size and gene ratios, the ratio of DEGs to total genes within the specified GO terms, are displayed on the x-axis. Gene-concept networks of top 5 enriched GO terms. Colored connections depict GO terms, nodes sizes indicate gene counts within the specified GO terms and fold changes in the KS signature cluster compared to the other clusters are indicated by color. (**E)** Venn diagram of DEGs identified in the KS signature cluster compared to other clusters, KSHV infected spots compared to uninfected spots, and two published bulk RNA sequencing datasets of KS tissue compared to normal skin (*34, 35*). Numbers of total DEGs per comparison and overlapping DEGs are provided.

**Supplementary Fig. 4**

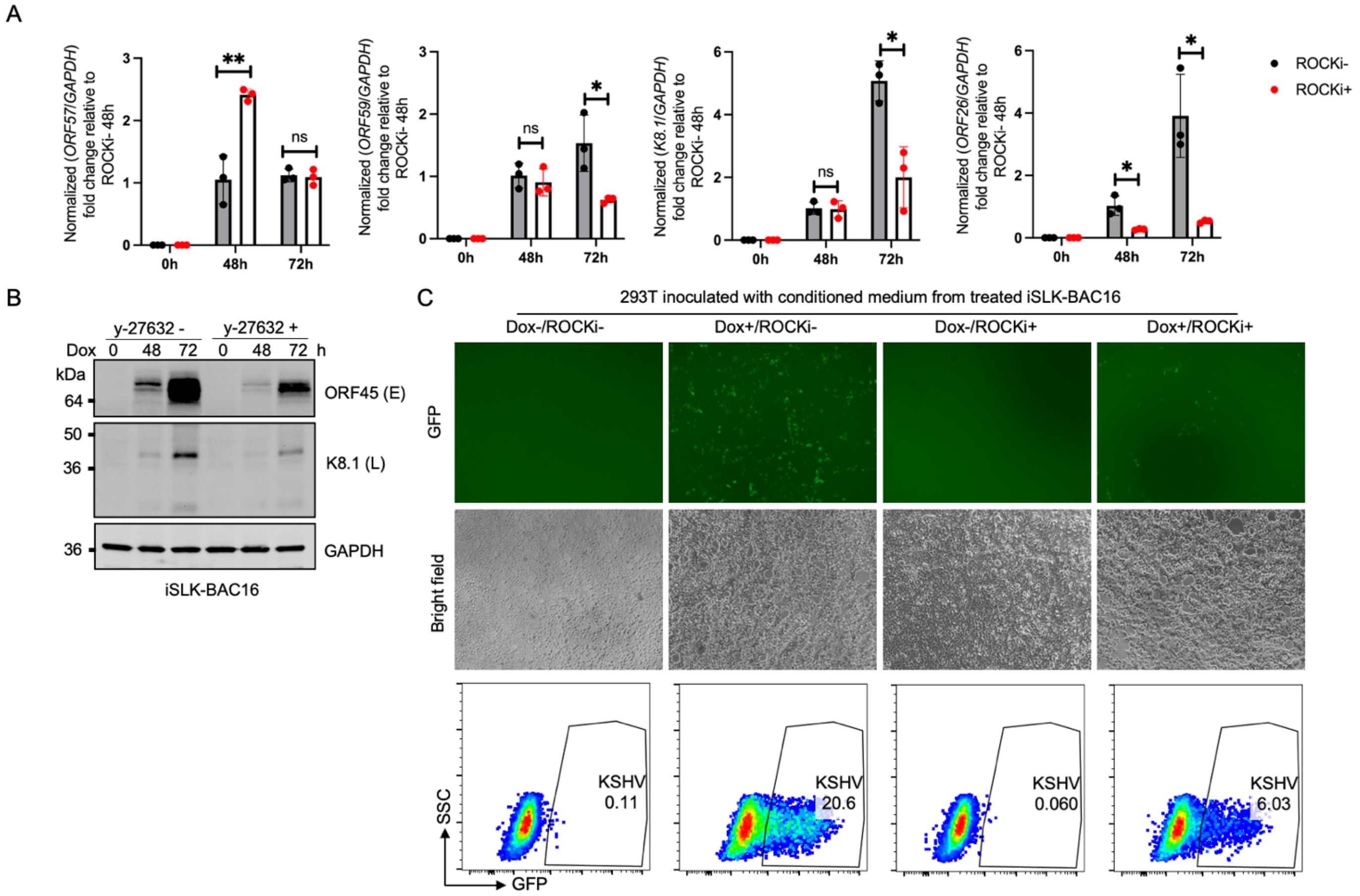

**Supplementary Fig. 4. KSHV lytic reactivation is impaired upon inhibition of Rho kinases.** iSLK-BAC16 cells were treated with 10 μM of Y-27632 for 2 hrs, then treated with 5 μg/ml of dox to induce KSHV reactivation. (**A-B**) Total RNA and cell lysates from treated iSLK-BAC16 cells were harvested at indicated timepoints post dox-induction for qRT-PCR to quantify KSHV transcripts (A) and immunoblotting for KSHV proteins (B). **(C)** Conditioned medium from iSLK-BAC16 collected at 96 h post dox-induction was used to inoculate 293T cells and evaluated for GFP as an indication of KSHV infection by microscopy and flow cytometry 48 hpi.

#### Supplementary Fig. 5

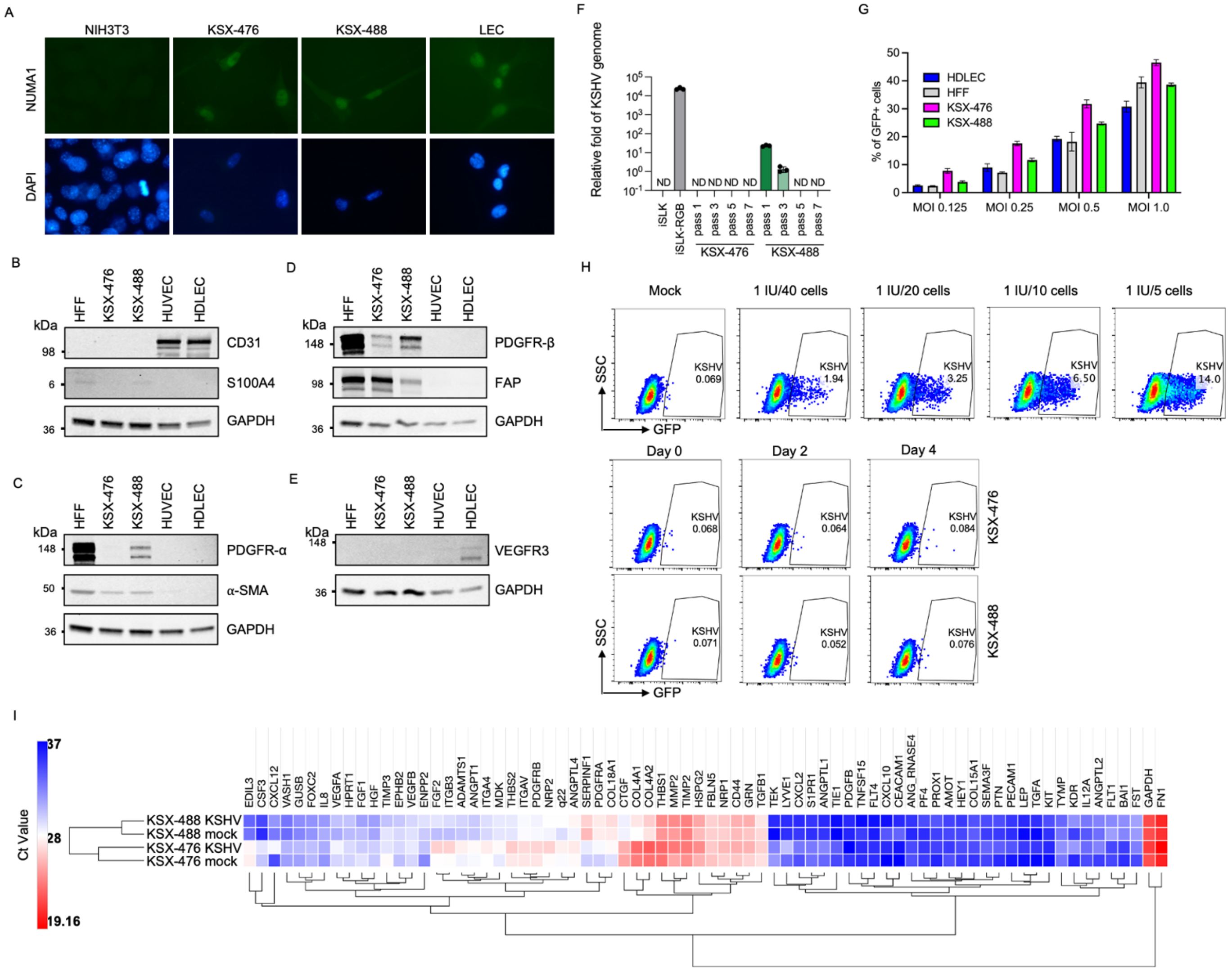

**Supplementary Fig. 5. KSX cells are human tumor associated fibroblast like cells that are KSHV negative but highly permissive to viral entry.**

**(A)** Mouse NIH3T3, KSXs and human LEC were fixed and stained with antibody against human cell marker NUMA1. (**B-E)** Cell lysates harvested from KSXs, HFF and endothelial cells were used for detecting markers of tumor associated fibroblast and endothelial cells by immunoblot as described in Fig. 7B. These blots from Fig. 7B are paired with GAPDH from the same blot as a loading control. **(F)** Quantitative PCR for the KSHV genome in total DNA extracted from KSXs at different passages. **(G)** 5 X $10^4$ LEC, HFF and KSXs were challenged with indicated infectious units of KSHV-BAC16, cells were harvest at 48 hpi for FACS to measure the percentage of GFP+ cells. **(H)** Conditioned media collected from mock and KSHV infected KSXs at indicated time points were used to incubate with 293T cells, and cells were harvested at 48 hpi for to detect GFP+ cells by FACS. KSHV-BAC16 virus was included in parallel as a control. **(I)** cDNAs harvested from mock KSX-476, KSHV infected KSX-476, mock KSX-488 or KSHV infected KSX-488 at 2 dpi were applied to qRT-PCR on four TaqMan™ Array Human Angiogenesis plates separately. 92 of target genes previously determined to be involved in angiogenesis and 4 housekeeping genes were examined for the Ct value with equal amount of cDNA loaded in each well. The heatmap generated with online software (https://software.broadinstitute.org/morpheus/) indicated the relative expression level of selected genes across 4 samples in the unit of Ct value.

**Supplementary Fig. 6**

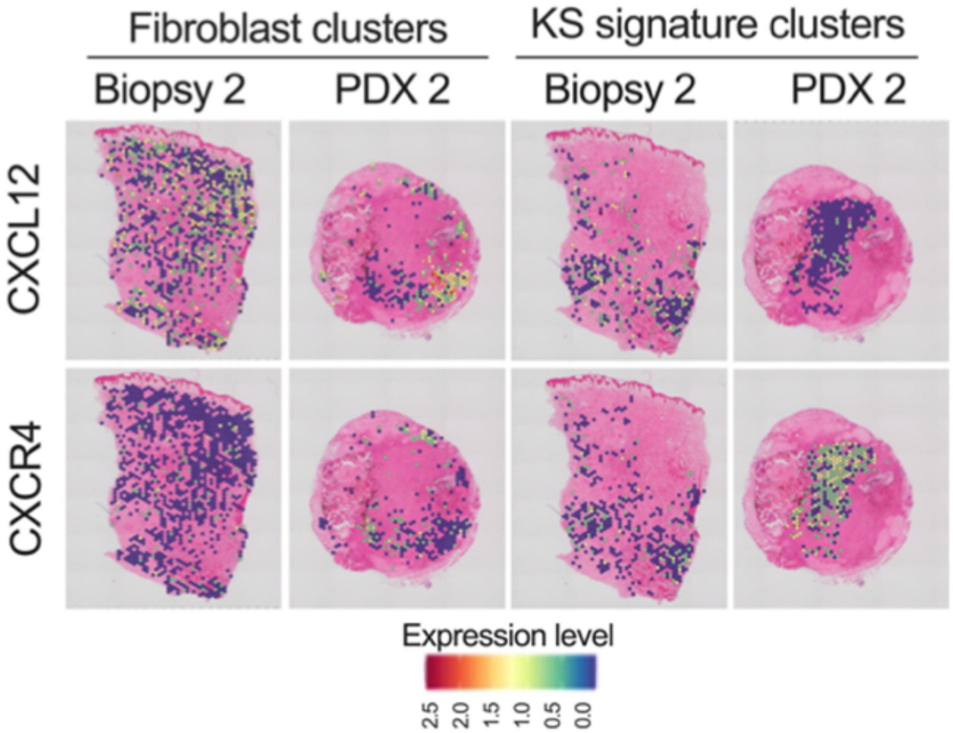


**Supplementary Fig. 6. Spatial distribution of CXCL12 and CXCR4.** SCT normalized expression levels of CXCL12 (upper) and CXCR4 (lower) SCT normalized expression levels in fibroblast and KS signature clusters in biopsy/PDX pair 2, as described for Fig. 7G.
